# Transduction of the Geomagnetic Field as Evidenced from Alpha-band Activity in the Human Brain

**DOI:** 10.1101/448449

**Authors:** Connie X. Wang, Isaac A. Hilburn, Daw-An Wu, Yuki Mizuhara, Christopher P. Cousté, Jacob N. H. Abrahams, Sam E. Bernstein, Ayumu Matani, Shinsuke Shimojo, Joseph L. Kirschvink

## Abstract

Magnetoreception, the perception of the geomagnetic field, is a sensory modality well-established across all major groups of vertebrates and some invertebrates, but its presence in humans has been tested rarely, yielding inconclusive results. We report here a strong, specific human brain response to ecologically-relevant rotations of Earth-strength magnetic fields. Following geomagnetic stimulation, a drop in amplitude of EEG alpha oscillations (8-13 Hz) occurred in a repeatable manner. Termed alpha event-related desynchronization (alpha-ERD), such a response is associated with sensory and cognitive processing of external stimuli. Biophysical tests showed that the neural response was sensitive to the dynamic components and axial alignment of the field but also to the static components and polarity of the field. This pattern of results implicates ferromagnetism as the biophysical basis for the sensory transduction and provides a basis to start the behavioral exploration of human magnetoreception.

## Introduction

Magnetoreception is a well-known sensory modality in bacteria (Frankel & Blakemore, 1980), protozoans (Bazylinski, Schlezinger, Howes, Frankel, & Epstein, 2000) and a variety of animals (Johnsen & Lohmann, 2008; Walker, Dennis, & Kirschvink, 2002; R. Wiltschko & W. Wiltschko, 1995), but whether humans have this ancient sensory system has never been conclusively established. Behavioral results suggesting that geomagnetic fields influence human orientation during displacement experiments (Baker, 1980, 1982, 1987) were not replicated (Able & Gergits, 1985; Gould & Able, 1981; Westby & Partridge, 1986). Attempts to detect human brain responses using electroencephalography (EEG) were limited by computational methods of the time (Sastre, Graham, Cook, Gerkovich, & Gailey, 2002). Twenty to thirty years after these previous flurries of research, the question of human magnetoreception remains unanswered.

In the meantime, there have been major advances in our understanding of animal geomagnetic sensory systems. An ever-expanding list of experiments on magnetically-sensitive organisms has revealed physiologically-relevant stimuli as well as environmental factors that may interfere with magnetosensory processing (Lohmann, Cain, Dodge, & Lohmann, 2001; Walker et al., 2002; R. Wiltschko & W. Wiltschko, 1995). Animal findings provide a potential feature space for exploring human magnetoreception – the physical parameters and coordinate frames to be manipulated in human testing (J. Kirschvink, Padmanabha, Boyce, & Oglesby, 1997; W. Wiltschko, 1972). In animals, geomagnetic navigation is thought to involve both a compass and map response (Kramer, 1953). The compass response simply uses the geomagnetic field as an indicator to orient the animal relative to the local magnetic north/south direction (Lohmann et al., 2001; R. Wiltschko & W. Wiltschko, 1995). The magnetic map is a more complex response involving various components of field intensity and direction; direction is further subdivided into inclination (vertical angle from the horizontal plane; the North-seeking vector of the geomagnetic field dips downwards in the Northern Hemisphere) and declination (clockwise angle of the horizontal component from Geographic North, as in a man-made compass). Notably, magnetosensory responses tend to shut down altogether in the presence of anomalies (e.g. sunspot activity or local geomagnetic irregularities) that cause the local magnetic field to deviate significantly from typical ambient values (Martin & Lindauer, 1977; W. Wiltschko, 1972), an adaptation that is thought to guard against navigational errors. These results indicate that geomagnetic cues are subject to complex neural processing, as in most other sensory systems.

Physiological studies have flagged the ophthalmic branch of the trigeminal system (and equivalents) in fish (Walker et al., 1997), birds (Beason & Semm, 1996; Elbers, Bulte, Bairlein, Mouritsen, & Heyers, 2017; Mora, Davison, Wild, & Walker, 2004; Semm & Beason, 1990) and rodents (Wegner, Begall, & Burda, 2006) as a conduit of magnetic sensory information to the brain. In humans, the trigeminal system includes many autonomic, visceral and proprioceptive functions that lie outside conscious awareness (Fillmore & Seifert, 2015; Saper, 2002). For example, the ophthalmic branch contains parasympathetic nerve fibers and carries signals of extraocular proprioception, which do not reach conscious awareness (Liu, 2005).

If the physiological components of a magnetosensory system have been passed from animals to humans, then their function may be either subconscious or only weakly available to conscious perception. Behavioral experiments could be easily confounded by cognitive factors such as attention, memory and volition, making the results weak or difficult to replicate at the group or individual levels. Since brain activity underlies all behavior, we chose a more direct electrophysiological approach to test for the transduction of geomagnetic fields in humans.

## Materials and Methods

We constructed an isolated, radiofrequency-shielded chamber wrapped with three nested sets of orthogonal square coils, using the four-coil design of Merritt *et al.* (Merritt, Purcell, & Stroink, 1983) for high central field uniformity (Fig. 1, and in the section on *Extended Materials and Methods* below). Each coil contained two matched sets of windings to allow operation in Active or Sham mode. Current ran in series through the two windings to ensure matched amplitudes. In Active mode, currents in paired windings were parallel, leading to summation of generated magnetic fields. In Sham mode, currents ran antiparallel, yielding no measurable external field, but with similar ohmic heating and magnetomechanical effects as in Active mode (J.L Kirschvink, 1992). Active and Sham modes were toggled by manual switches in the distant control room, leaving computer and amplifier settings unchanged. Coils were housed within an acoustically-attenuated, grounded Faraday cage with aluminum panels forming the walls, floor and ceiling. Participants sat upright in a wooden chair on a platform electrically isolated from the coil system with their heads positioned near the center of the uniform field region and their eyes closed in total darkness. (Light levels within the experimental chamber during experimental runs were measured using a Konica-Minolta CS-100A luminance meter, which gave readings of zero, e.g. below 0.01 ± 2% cd/m^2^.) The magnetic field inside the experimental chamber was monitored by a three-axis Applied Physics Systems^™^ 520A fluxgate magnetometer. EEG was continuously recorded from 64 electrodes using a BioSemi^™^ ActiveTwo system with electrode positions coded in the International 10-20 System (e.g. Fz, CPz, etc.). Inside the cage, the battery-powered digital conversion unit relayed data over a non-conductive, optical fiber cable to a remote control room, ∼20 meters away, where all power supplies, computers and monitoring equipment were located.

**Fig 1.**
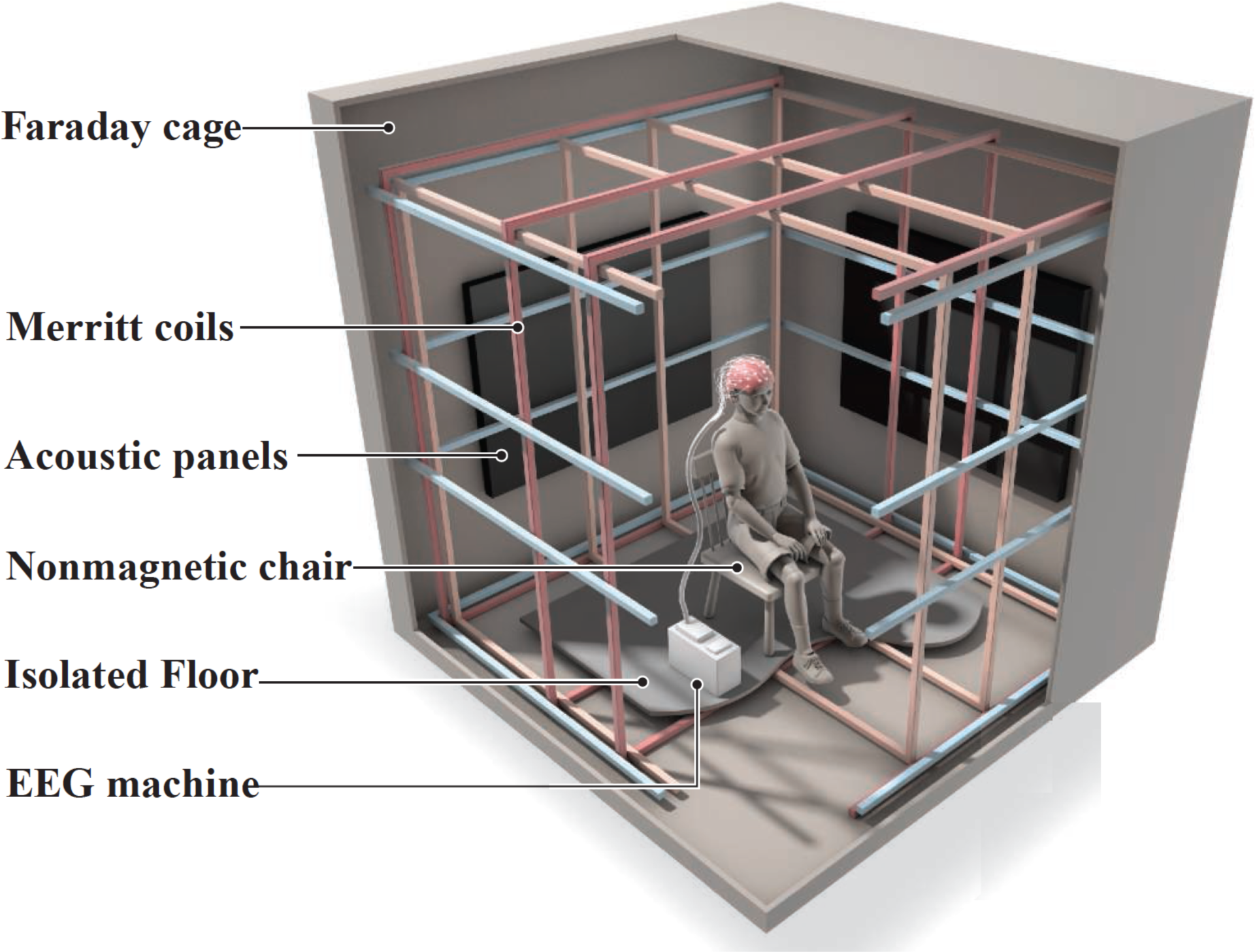
Schematic illustration of the experimental setup. The ∼1 mm thick aluminum panels of the electrically-grounded Faraday shielding provides an electromagnetically “quiet” environment. Three orthogonal sets of square coils ∼2 m on edge, following the design of Merritt *et al.* (Merritt et al., 1983), allow the ambient geomagnetic field to be altered around the participant’s head with high spatial uniformity; double-wrapping provides an active-sham for blinding of experimental conditions (J.L Kirschvink, 1992). Acoustic panels on the wall help reduce external noise from the building air ventilation system as well as internal noise due to echoing. A non-magnetic chair is supported on an elevated wooden base isolated from direct contact with the magnetic coils. The battery-powered EEG is located on a stool behind the participant and communicates with the recording computer via an optical fiber cable to a control room ∼20 m away. Additional details are available in the *Extended Materials and Methods* section, and Fig. 5 below. This diagram was modified from the figure “Center of attraction”, by C. Bickel (Hand, 2016), with permission.

A ∼1 hour EEG session consisted of multiple ∼7 minute experimental runs. In each run of 100+ trials, magnetic field direction rotated repeatedly between two preset orientations with field intensity held nearly constant at the ambient lab value (∼35 μT). In SWEEP trials, the magnetic field started in one orientation then rotated smoothly over 100 milliseconds to the other orientation. As a control condition, FIXED trials with no magnetic field rotation were interspersed amongst SWEEP trials according to pseudorandom sequences generated by software. Trials were separated in time by 2-3 seconds. The experimental chamber was dark, quiet and isolated from the control room during runs. Participants were blind to Active vs. Sham mode, trial sequence and trial timing. During sessions, auditory tones signaled the beginning and end of experiment runs, and experimenters only communicated with participants once or twice per session between active runs to update the participant on the number of runs remaining. When time allowed, Sham runs were matched to Active runs using the same software settings. Active and Sham runs were programmatically identical, differing only in the position of hardware switches that directed current to run parallel or antiparallel through paired loops. Sham runs served as an additional control for non-magnetic sensory confounds, such as sub-aural stimuli or mechanical oscillations from the coil system. (Note that experimental variables differing *between* runs are denoted in camel case as in DecDn, DecUp, Active, Sham, etc., whereas variables that change *within* runs are designated in all capitals like FIXED, SWEEP, CCW, CW, UP, DN, etc.). In Active runs, an electromagnetic induction artifact occurred as a 10-20 microvolt fluctuation in the EEG signal during the 100 ms magnetic field rotation. This induction artifact is similar to that observed in electrophysiological recordings from trout whenever magnetic field direction or intensity was suddenly changed in a square wave pattern (Walker et al., 1997). Strong induced artifacts also occur in EEG recordings during transcranial magnetic stimulation (TMS) (Veniero, Bortoletto, & Miniussi, 2009). In all cases, the artifact can only be induced in the presence of time-varying magnetic fields and disappears once the magnetic field stabilizes (∂B/∂t=0). In our experiments, EEG data following the 100 ms field rotation interval were not subject to effects from the induction artifact. Furthermore, the induction artifact is phase-locked like an event-related potential and does not appear in analyses of non-phase-locked power, which we used in all subsequent statistical tests. Further discussion of electrical induction is in section 4 of *Extended Materials and Methods,* below.

Fig. 2 shows the magnetic field rotations used. In inclination (Inc) experiments (Fig. 2A), declination direction was fixed to North (0° declination in our coordinate system), and participants sat facing North. Rotation of the field vector from downwards to upwards was designated as an ‘Inc.UP.N’ trial and the return sweep as ‘Inc.DN.N’, with UP/DN indicating the direction of field rotation. In declination (Dec) experiments (Fig 2B, 2C), we held inclination (and hence the vertical component of the field vector) constant, while rotating the horizontal component clockwise or counterclockwise to vary the declination. For trials with downwards inclination (as in the Northern Hemisphere), field rotations swept the horizontal component 90° CW or CCW between Northeast and Northwest, designated as ‘DecDn.CW.N’ or ‘DecDn.CCW.N’, respectively, with ‘.N’ indicating a Northerly direction. To test biophysical hypotheses of magnetoreception as discussed below, we conducted additional declination rotation experiments with static, upwards inclination. As shown in Fig. 2B, rotating an upwards-directed field vector between SE and SW (‘DecUp.CW.S’ and ‘DecUp.CCW.S’) antiparallel to the downwards-directed rotations provides tests of the quantum compass biophysical model, while sweeping an upwards vector between NE and NW (‘DecUp.CW.N’ and ‘DecUp.CCW.N’) provides a general test for electrical induction (Fig. 2C).

**Fig 2.**
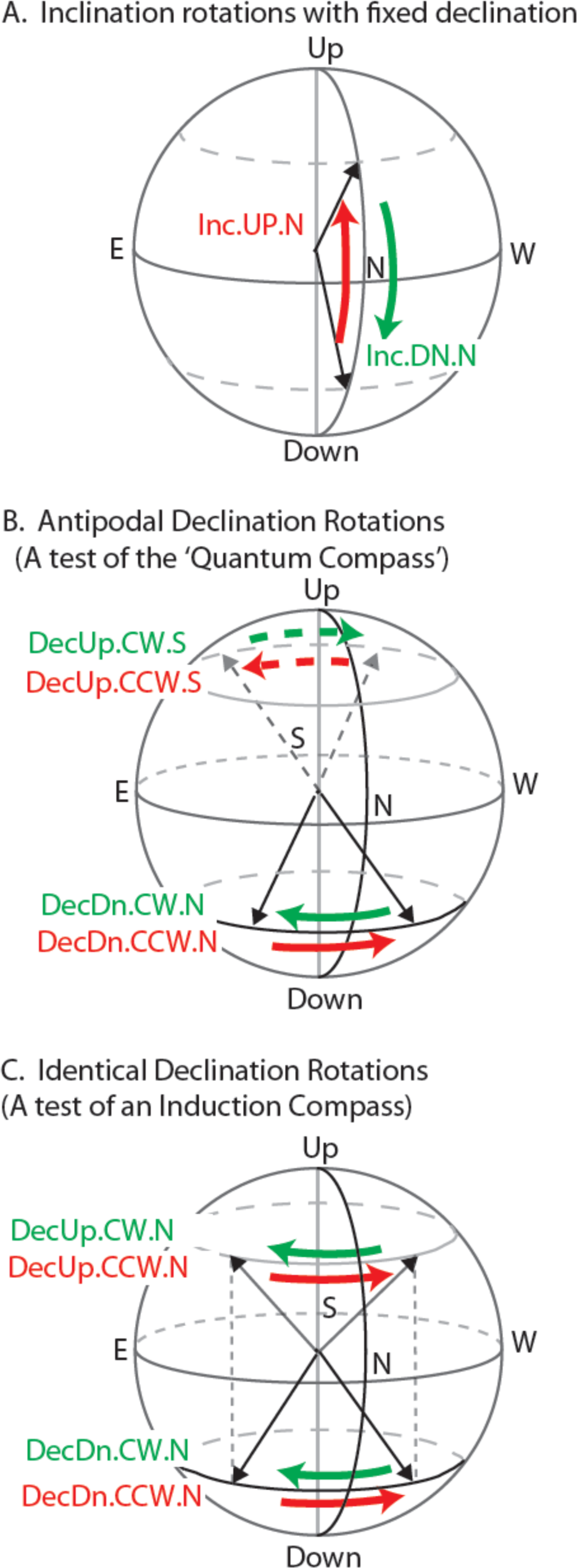
Magnetic field rotations used in these experiments. In the first ∼100 ms of each experimental trial, the magnetic field vector was either: 1) rotated from the first preset orientation to the second (SWEEP), 2) rotated from the second preset orientation to the first (also SWEEP), or 3) left unchanged (FIXED). In all experimental trials, the field intensity was held constant at the ambient lab value (∼35 uT). For declination rotations, the horizontal rotation angle was +90 degrees or −90 degrees. For inclination rotations, the vertical rotation angle was either +120 degrees / −120 degrees, or +150 degrees / −150 degrees, depending on the particular inclination rotation experiment. (A) Inclination rotations between ±60° or ±75°. The magnetic field vector rotates from downwards to upwards (Inc.UP.N, red) and vice versa (Inc.DN.N, green), with declination steady at North (0°). (B) Declination rotations used in main assay (solid arrows) and vector opposite rotations used to test the quantum compass hypothesis (dashed arrows). In the main assay, the magnetic field rotated between NE (45°) and NW (315°) with inclination held downwards (+60° or +75°) as in the Northern Hemisphere (DecDn.CW.N and DecDn.CCW.N); vector opposites with upwards inclination (−60° or −75°) and declination rotations between SE (135°) and SW (225°) are shown with dashed arrows (DecUp.CW.S and DecUp.CCW.S). (C) Identical declination rotations, with static but opposite vertical components, used to test the electrical induction hypothesis. The magnetic field was shifted in the Northerly direction between NE (45°) and NW (315°) with inclination held downwards (+75°, DecDn.CW.N and DecDn.CCW.N) or upwards (−75°, DecUp.CW.S and DecUp.CCW.S). The two dotted vertical lines indicate that the rotations started at the same declination values. In both (B) and (C), counterclockwise rotations (viewed from above) are shown in red, clockwise in green.

During magnetic field rotations, EEG was recorded from participants in the eyes-closed resting state. Auditory cues marked the beginning and end of each ∼7 minute run, but participants were not informed of run mode, trial sequence or stimulus timing. EEG was sampled at 512 Hz from 64 electrodes arrayed in the standard International 10-20 positions using a Biosemi^™^ ActiveTwo system. The experimental protocol was approved by the Caltech Institutional Review Board (IRB), and all participants gave written informed consent.

We used conventional methods of time/frequency decomposition (Morlet wavelet convolution) to compute post-stimulus power changes relative to a pre-stimulus baseline interval (−500 to −250 ms) over a 1-100 Hz frequency range. We focused on non-phase-locked power by subtracting the event-related potential in each condition from each trial of that condition prior to time/frequency decomposition. This is a well-known procedure for isolating non-phase-locked power and is useful for excluding the artifact from subsequent analyses (Cohen, 2014). Following the identification of alpha band activity as a point of interest (detailed in Results), the following procedure was adopted to isolate alpha activity in individuals. To compensate for known individual differences in peak resting alpha frequency (8 to 12 Hz in our participant pool) and in the timing of alpha wave responses following sensory stimulation, we identified individualized power change profiles using an automated search over an extended alpha band of 6-14 Hz, 0-2 s post-stimulus. For each participant, power changes at electrode Fz were averaged over all trials, regardless of condition, to produce a single time/frequency map. In this cross-conditional average, the most negative time-frequency point was set as the location of the participant’s characteristic alpha-ERD. A window of 250 ms and 5 Hz bandwidth was automatically centered as nearly as possible on that point within the constraints of the overall search range. These time/frequency parameters were chosen based on typical alpha-ERD durations and bandwidths. Alpha power activity in each individualized window was used to test for significant differences between conditions. For each condition, power changes were averaged separately within the window, with trials subsampled and bootstrapped to equalize trial numbers across conditions. Outlier trials with extreme values of alpha power (typically caused by movement artifacts or brief bursts of alpha activity in an otherwise low-amplitude signal) in either the pre-or post-stimulus intervals were removed by an automated algorithm prior to averaging, according to a threshold of 1.5X the interquartile range of log alpha power across all trials. Further details are provided in sections 1-5 of *Extended Materials and Methods,* below.

## Results

In initial observations, several participants (residing in the Northern Hemisphere) displayed striking patterns of neural activity following magnetic stimulation, with strong decreases in EEG alpha power in response to two particular field rotations: (1) Inclination SWEEP trials (Inc.UP.N and Inc.DN.N), in which the magnetic vector rotated either down or up (e.g. rotating a downwards pointed field vector up to an upwards pointed vector, or vice versa; Fig. 2A red and green arrows), and (2) DecDn.CCW.N trials, in which magnetic field declination rotated counterclockwise while inclination was held downwards (as in the Northern Hemisphere; Fig 2B, solid red arrow). Alpha power began to drop from pre-stimulus baseline levels as early as ∼100 ms after magnetic stimulation, decreasing by as much as ∼50% over several hundred milliseconds, then recovering to baseline by ∼1 s post-stimulus; this is visualized by the deep blue color on the time-frequency power maps (Fig. 3). Scalp topography was bilateral and widespread, centered over frontal/central electrodes, including midline frontal electrode Fz when referenced to CPz. Fig. 3A shows the whole-brain response pattern to inclination sweeps and control trials (Inc.SWEEP.N and Inc.FIXED.N) of one of the responsive participants, with the alpha-ERD exhibited in the SWEEP but not FIXED trials. Similarly, Fig. 3B and 3C show the declination responses of a different participant on two separate runs (labeled Runs #1 and #2) six months apart. Response timing, bandwidth and topography of the alpha-ERD in the CCW sweeps, with negative FIXED controls, were replicated across runs, indicating a repeatable signature of magnetosensory processing in humans. After experimental sessions, participants reported that they could not discern when or if any magnetic field changes had occurred.

**Fig 3.**
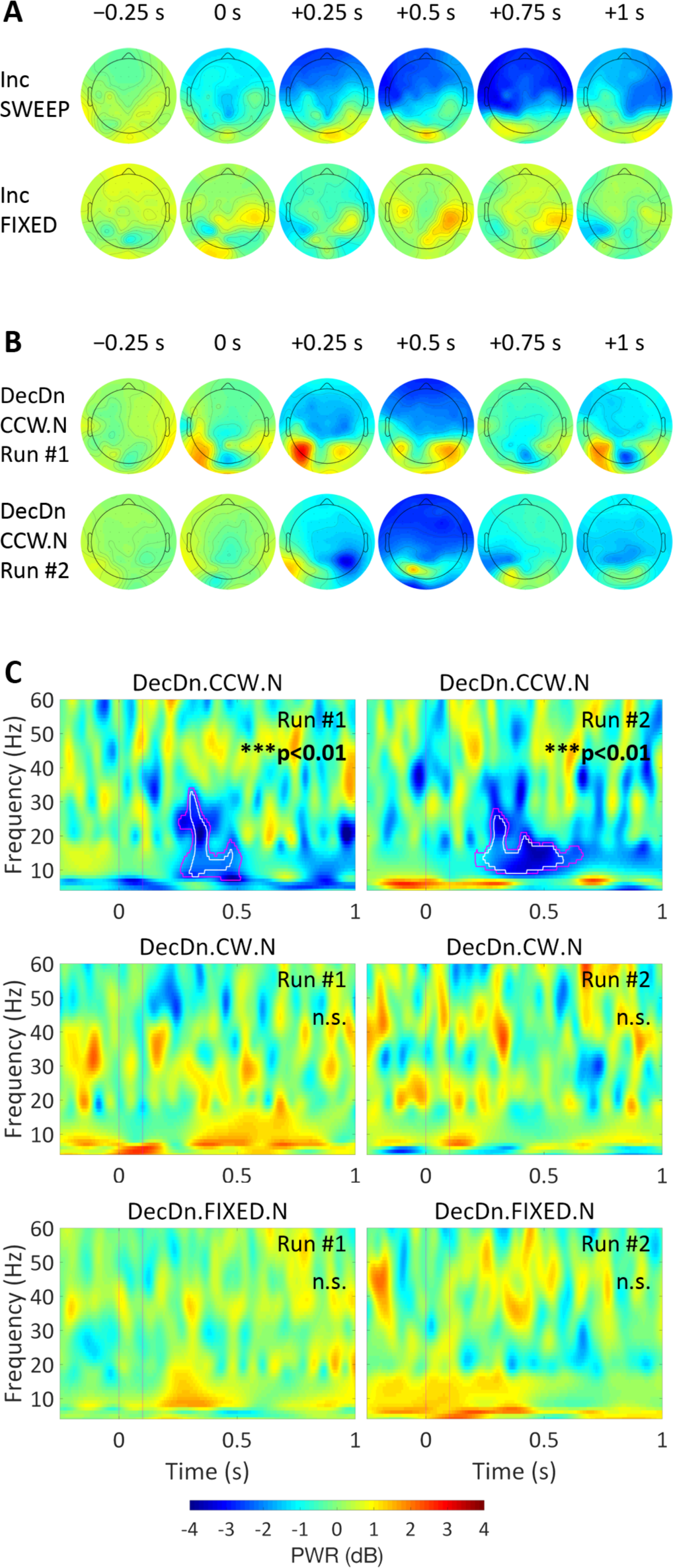
Alpha-ERD as a neural response to magnetic field rotation. Post-stimulus power changes (dB) from a pre-stimulus baseline (−500 to −250 ms) plotted according to the ±4 dB color bar at bottom. **(A)** Scalp topography of the alpha-ERD response in an inclination experiment, showing alpha power at select time points before and after field rotation at 0 s. Alpha-ERD (deep blue) was observed in SWEEP (top row), but not FIXED (bottom row), trials. **(B)** Scalp topography of the alpha-ERD response for two runs of the declination experiment, tested 6 months apart in a different strongly-responding participant. DecDn.CCW.N condition is shown. In both runs, the response peaked around +500 ms post-stimulus and was widespread over frontal/central electrodes, demonstrating a stable and reproducible response pattern. **(C)** Time-frequency maps at electrode Fz for the same runs shown in (B). Pink vertical lines indicate the 0-100 ms field rotation interval. Pink/white outlines indicate significant alpha-ERD at the p<0.05 and p<0.01 statistical thresholds, respectively. Separate runs shown side by side. Significant alpha-ERD was observed following downwards-directed counterclockwise rotations (outlines in top row), with no other power changes reaching significance. Significant power changes appear with similar timing and bandwidth, while activity outside the alpha-ERD response, and activity in other conditions is inconsistent across runs.

The alpha rhythm is the dominant human brain oscillation in the resting state when a person is not processing any specific stimulus or performing any specific task (Klimesch, 1999). Neurons engaged in this internal rhythm produce 8-13 Hz alpha waves that are measurable by EEG. Individuals vary widely in the amplitude of the resting alpha rhythm. When an external stimulus is suddenly introduced and processed by the brain, the alpha rhythm generally decreases in amplitude compared with a pre-stimulus baseline. (Hartmann, Schlee, & Weisz, 2012; Klimesch, 1999; Pfurtscheller, Neuper, & Mohl, 1994). This EEG phenomenon, termed alpha event-related desynchronization (alpha-ERD), has been widely observed during perceptual and cognitive processing across visual, auditory and somatosensory modalities (Peng, Hu, Zhang, & Hu, 2012). Alpha-ERD may reflect the recruitment of neurons for processing incoming sensory information and is thus a generalized signature for a shift of neuronal activity from the internal resting rhythm to external engagement with sensory or task-related processing (Pfurtscheller & Lopes da Silva, 1999). Individuals also vary in the strength of alpha-ERD; those with high resting-state or pre-stimulus alpha power tend to show strong alpha-ERDs following sensory stimulation, while those with low alpha power have little or no response in the alpha band (Klimesch, 1999).

Based on early observations, we formed the hypothesis that sensory transduction of geomagnetic stimuli could be detectable as alpha-ERD in response to field rotations – e.g. the magnetic field rotation would be the external stimulus, and the alpha-ERD would be the signature of the brain beginning to process sensory data from this stimulus. This hypothesis was tested at the group level in data collected from 29 participants in the inclination rotation conditions (Fig 2A) and 26 participants in the declination rotation conditions (Fig. 2B, solid arrows).

For inclination experiments, we collected data from matched Active and Sham runs (N=29 of 34; 5 participants were excluded due to time limits that prevented the collection of sham data). We tested for the effects of inclination rotation (SWEEP vs. FIXED) and magnetic stimulation (Active vs. Sham) using a two-way repeated-measures ANOVA. We found a significant interaction of inclination rotation and magnetic stimulation (p<0.05). Post-hoc comparison of the four experimental conditions (Active-SWEEP, Active-FIXED, Sham-SWEEP, Sham-FIXED) revealed significant differences between Active-SWEEP and all other conditions (p<0.05). Downwards/upwards rotations of magnetic field inclination produced an alpha-ERD ∼2X greater than background fluctuations in the FIXED control condition and all the Sham conditions. Results are summarized in Table 1 and Fig. 4A.

**Fig 4.**
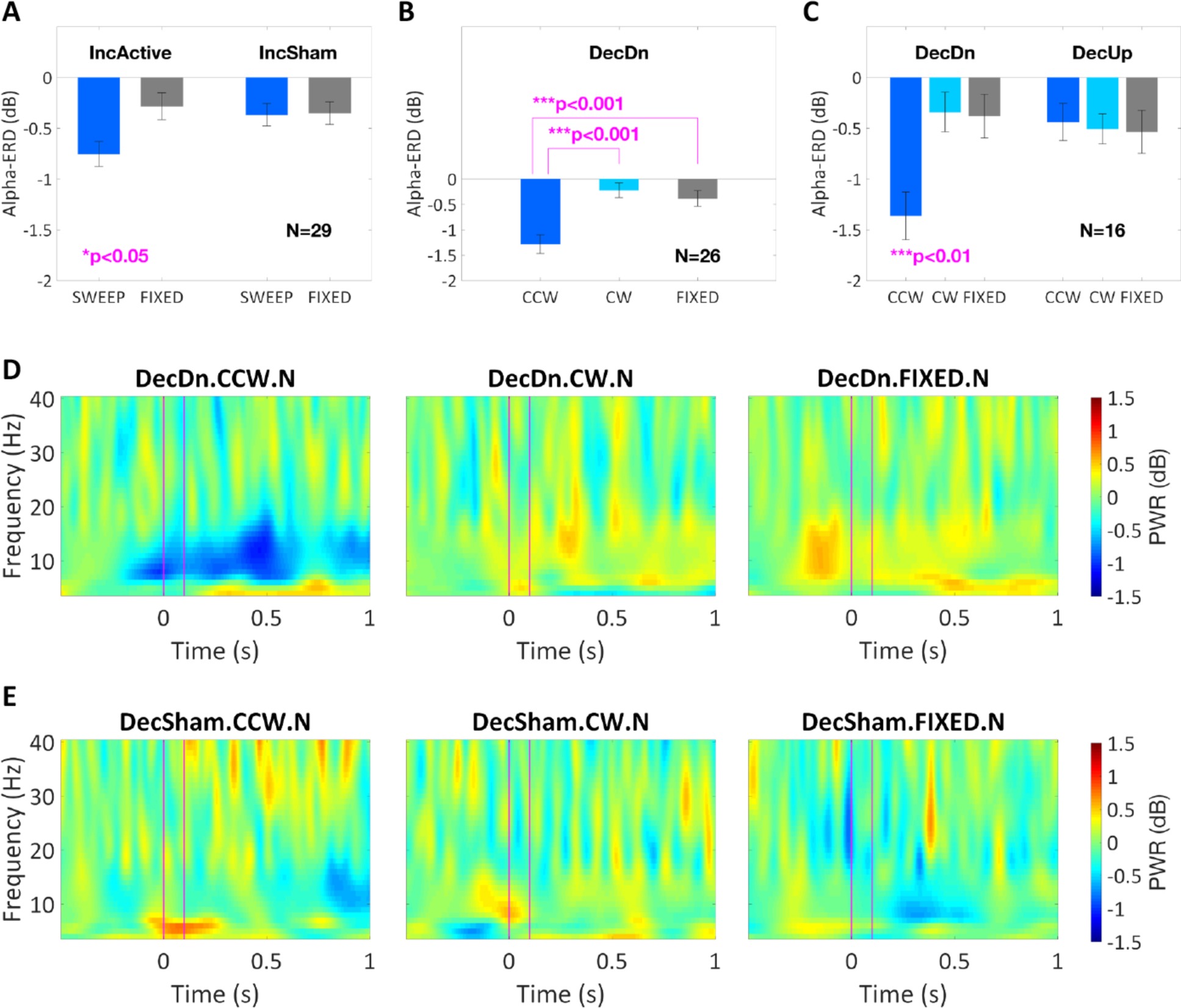
Group results from repeated-measures ANOVA for the effects of geomagnetic stimulation on post-stimulus alpha power. (A) Average alpha-ERD (dB) at electrode Fz in the SWEEP and FIXED conditions of inclination experiments run in Active or Sham mode. Two-way ANOVA showed an interaction (p<0.05, N=29) of inclination rotation (SWEEP vs. FIXED) and magnetic stimulation (Active vs. Sham). According to post-hoc testing, only inclination sweeps in Active mode produced alpha-ERD above background fluctuations in FIXED trials (p<0.01) or Sham mode (p<0.05). (B) Average alpha-ERD (dB) at electrode Fz in the declination experiment with inclination held downwards (DecDn). One-way ANOVA showed a significant main effect of declination rotation (p<0.001, N=26). The downwards-directed counterclockwise rotation (DecDn.CCW.N) produced significantly different effects from both the corresponding clockwise rotation (DecDn.CW.N, p<0.001) and the FIXED control condition (DecDn.FIXED.N, p<0.001). (C) Comparison of the declination rotations with inclination held downwards (DecDn) or upwards (DecUp) in a subset (N=16 of 26) of participants run in both experiments. Two-way ANOVA showed a significant interaction (p<0.01) of declination rotation (CCW vs. CW vs. FIXED) and inclination direction (Dn vs. Up). Post-hoc testing showed significant differences (p<0.01) between the DecDn.CCW.N condition and every other condition, none of which were distinct from any other. This is a direct test and rejection of the quantum compass hypothesis. (D) Grand average of time-frequency power changes across the 26 participants in the DecDn experiment from (B). Pink vertical lines indicate the 0-100 ms field rotation interval. A post-stimulus drop in alpha power was observed only following the downwards-directed counterclockwise rotation (left panel). Wider spread of desychronization reflects inter-individual variation. Convolution involved in time/frequency analyses causes the early responses of a few participants to appear spread into the pre-stimulus interval. (E) Grand average of time-frequency power changes across the 18 participants with sham data in the declination experiments; no significant power changes were observed.

**Table 1.** Group results from repeated-measures ANOVA for the effects of magnetic field rotation on post-stimulus alpha power. ANOVA #1 shows a significant interaction of inclination rotation (SWEEP vs. FIXED) and magnetic stimulation (Active vs. Sham) in the inclination experiments. Based on post-hoc testing, alpha-ERD was significantly greater in SWEEP trials in Active mode, compared with all other conditions (p<0.05). ANOVA #2 shows a significant main effect of declination rotation when inclination is static and downwards as in the Northern Hemisphere. Alpha-ERD was significantly greater following counterclockwise rotations (p<0.001). ANOVA #3 shows a significant interaction of declination rotation and inclination direction in declination experiments designed to test the “Quantum Compass” mechanism of magnetoreception. A significant alpha-ERD difference (p<0.01) between counterclockwise down (DecDn.CCW.N) and counterclockwise up (DecUp.CCW.S) argues against this hypothesis in humans.

| Group Results for Effects of Magnetic Field Rotation on Post-Stimulus Alpha Power |  |  |  |
| --- | --- | --- | --- |
| ANOVA #1: Inclination Rotation x Magnetic Stimulation (N=29) |  |  |  |
| Two-Way Repeated Measures ANOVA | F | p | $\eta_p^2$ |
| Main Effect of Inclination Rotation<br>(SWEEP vs. FIXED) | 3.26 | 0.08 | 0.19 |
| Main Effect of Magnetic Stimulation<br>(Active vs. Sham) | 2.47 | 0.13 | 0.09 |
| Inclination Rotation x Magnetic Stimulation<br>(Interaction) | 5.67 | **0.02\*** | 0.17 |
| ANOVA #2: Declination Rotation (N=26) |  |  |  |
| One-Way Repeated Measures ANOVA | F | p | $\eta_p^2$ |
| Main Effect of Declination Rotation<br>(CCW vs. CW vs. FIXED) | 13.09 | **0.00003\*\*\*** | 0.34 |
| ANOVA #3: Declination Rotation x Inclination Direction (N=16) |  |  |  |
| Two-Way Repeated Measures ANOVA | F | p | $\eta_p^2$ |
| Main Effect of Declination Rotation<br>(CCW vs. CW vs. FIXED) | 3.77 | **0.03\*** | 0.24 |
| Main Effect of Inclination Direction<br>(Dn vs. Up) | 0.89 | 0.36 | 0.06 |
| Declination Rotation x Inclination Direction<br>(Interaction) | 6.49 | **0.004\*\*\*** | 0.30 |

In declination experiments (Fig. 4B), we observed a strikingly asymmetric response to the clockwise (DecDn.CW.N) and counterclockwise (DecDn.CCW.N) rotations of a downwards-directed field sweeping between Northeast and Northwest. Alpha-ERD was ∼3X greater after counterclockwise than after clockwise rotations, the latter producing alpha power changes indistinguishable from background fluctuations in the FIXED control condition. Over the participant pool (N=26 of 26 who were run in this experiment), we ran a one-way repeated-measures ANOVA with three conditions (DecDn.CCW.N, DecDn.CW.N, and DecDn.FIXED.N) to find a highly significant effect of declination rotation (p<0.001) (Table 1). As indicated in Fig. 4B, the counterclockwise rotation elicited a significantly different response from both the clockwise rotation (p<0.001) and FIXED control (p<0.001). Fig. 4D shows the stimulus-locked grand average across all participants for each condition; an alpha-ERD is observed only for counterclockwise rotations of a downwards-directed field (left panel). Sham data were available for 18 of 26 participants in the declination experiments; no major changes in post-stimulus power were observed in any of the sham conditions (Fig. 4E).

The asymmetric declination response provided a starting point for evaluating potential mechanisms of magnetosensory transduction, particularly the quantum compass hypothesis, which has received much attention in recent years (Hore & Mouritsen, 2016; Ritz, Adem, & Schulten, 2000). Because the quantum compass cannot distinguish polarity, we conducted additional declination rotation experiments in which the fields were axially identical to those in the preceding DecDn experiments, except with reversed polarity (Fig. 2B; reversed polarity rotations shown as dashed arrows). In the additional DecUp conditions, Magnetic North pointed to Geographic South and up rather than Geographic North and down, and the upwards-directed field rotated clockwise (DecUp.CW.S) or counterclockwise (DecUp.CCW.S) between SE and SW. In later testing, we ran 16 participants in both the DecDn and DecUp experiments to determine the effects of declination rotation and inclination direction in a two-way repeated measures ANOVA with six conditions (DecDn.CCW.N, DecDn.CW.N, DecDn.FIXED.N, DecUp.CCW.S, DecUp.CW.S, and DecUp.FIXED.S). A significant interaction of declination rotation and inclination direction (p<0.01) was found (Fig. 4C and Table 1). DecDn.CCW.N was significantly different from all other conditions (p<0.01), none of which differed from any other. Thus, counterclockwise rotations of a downwards-directed field were processed differently in the human brain from the same rotations of a field of opposite polarity. These results contradict the quantum compass hypothesis, as explained below in Biophysical Mechanisms.

From previous EEG studies of alpha oscillations in human cognition, the strength of alpha-ERD is known to vary substantially across individuals (Klimesch, 1999; Klimesch, Doppelmayr, Russegger, Pachinger, & Schwaiger, 1998; Pfurtscheller et al., 1994). In agreement with this, we observed a wide range of alpha-ERD responses in our participants as well. Some participants showed large drops in alpha power up to ∼60% from pre-stimulus baseline, while others were unresponsive with little change in post-stimulus power at any frequency. Histograms of these responses are provided in Fig. 6-8 of *Extended Materials and Methods* below.

To confirm that the variability across the dataset was due to characteristic differences between individuals rather than general variability in the measurement or the phenomenon, we retested the strongly-responding participants to see if their responses were stable across sessions. Using permutation testing with false discovery rate (FDR) correction at the p<0.05 and p<0.01 statistical thresholds, we identified participants who exhibited alpha-ERD that reached significance at the individual level and tested them (N=4) again weeks or months later. An example of separate runs on the same participant is shown in Figs. 3B and 3C, and further data series are shown in the Fig 9 of *Extended Materials and Methods.* Each participant replicated their results with similar response tuning, timing and topography, providing greater confidence that the observed effect was specific for the magnetic stimulus in the brain of that individual. While the functional significance of these inter-individual differences is unclear, the identification of strongly responding individuals gives us the opportunity to conduct more focused tests directed at deriving the biophysical characteristics of the transduction mechanism.

### Biophysical Mechanisms

Three major biophysical transduction hypotheses have been considered extensively for magnetoreception in animals: (1) various forms of electrical induction (Kalmijn, 1981; Rosenblum, Jungerman, & Longfellow, 1985; Yeagley, 1947), (2) a chemical/quantum compass involving hyperfine interactions with a photoactive pigment (Schulten, 1982) like cryptochrome (Hore & Mouritsen, 2016; Ritz et al., 2000), and (3) specialized organelles based on biologically-precipitated magnetite similar to those in magnetotactic microorganisms (J.L. Kirschvink & Gould, 1981). We designed the declination experiments described above to test these hypotheses.

#### Electrical Induction

According to the Maxwell-Faraday law (∇ × **E** = −∂**B**/∂t), electrical induction depends only on the component of the magnetic field that is changing with time (∂**B**/∂t). In our declination experiments, this corresponds to the horizontal component that is being rotated. The vertical component is held constant and therefore does not contribute to electrical induction. Thus, we compared brain responses to two matched conditions, where the declination rotations were identical, but the static vertical components were opposite (Fig 2C). A transduction mechanism based in electrical induction would respond identically to these two conditions. Video 1 shows the alpha-ERD magnetosensory response of one strongly-responding individual to these two stimulus types. In the top row, the static component was pointing upwards, and in the bottom row, the static field was pointing downwards. In the DecDn.CCW.N condition (lower left panel), the alpha-ERD (deep blue patch) starts in the right parietal region almost immediately after magnetic stimulation and spreads over the scalp to most recording sites. This large, prolonged and significant bilateral desynchronization (p<0.01 at Fz) occurs only in this condition with only shorter, weaker and more localized background fluctuations in the other conditions (n.s. at Fz). No alpha-ERD was observed following any upwards-directed field rotation (DecUp.CCW.N and DecUp.CW.N, top left and middle panels), in contrast to the strong response in the DecDn.CCW.N condition. Looking across all of our data, none of our experiments (on participants from the Northern Hemisphere) produced alpha-ERD responses to rotations with a static vertical-upwards magnetic field (found naturally in the Southern Hemisphere).

These tests indicate that electrical induction mechanisms cannot account for the neural response. This analysis also rules out an electrical artifact of induced current loops from the scalp electrodes, as any current induced in the loops would also be identical across the matched runs. Our results are also consistent with many previous biophysical analyses, which argue that electrical induction would be a poor transduction mechanism for terrestrial animals, as the induced fields are too low to work reliably without large, specialized anatomical structures that would have been identified long ago (Rosenblum et al., 1985; Yeagley, 1947). Other potential confounding artifacts are discussed in sections 6 and 7 of the *Extended Materials and Methods,* below.

#### Quantum Compass

From basic physical principles, a transduction mechanism based on quantum effects can be sensitive to the axis of the geomagnetic field but not the polarity (Ritz et al., 2000; Schulten, 1982). In the most popular version of this theory, a photosensitive molecule like cryptochrome absorbs a blue photon, producing a pair of free radicals that can transition between a singlet and triplet state, with the transition frequency depending on the local magnetic field. The axis of the magnetic field – but not the polarity – could then be monitored by differential reaction rates from the singlet vs. triplet products. This polarity insensitivity, shared by all quantum-based magnetotransduction theories, is inconsistent with the group level test of the quantum compass presented above. The data (Table 1 and Figure 4C, dark blue bars) showed clearly distinct responses depending on polarity. We additionally verified this pattern of results at the individual level. Video 2 shows the alpha-ERD magnetosensory response in another strongly-responding individual. Only the DecDn.CCW.N rotation (lower left panel) yields a significant alpha-ERD (p<0.01 at Fz). Lack of a significant response in the axially identical DecUp.CCW.S condition indicates that the human magnetosensory response is sensitive to polarity. This means that a quantum compass-based mechanism cannot account for the alpha-ERD response we observe in humans.

### Response Selectivity

The selectivity of brain responses for specific magnetic field directions and rotations may be explained by tuning of neural activity to ecologically relevant values. Such tuning is well known in marine turtles in the central Atlantic Ocean, where small increases in the local geomagnetic inclination or intensity (that indicate the animals are drifting too far North and are approaching the Gulf Stream currents) trigger abrupt shifts in swimming direction, thereby preventing them from being washed away from their home in the Sargasso Sea (Light, Salmon, & Lohmann, 1993; Lohmann et al., 2001; Lohmann & Lohmann, 1996). Some migratory birds are also known to stop responding to the magnetic direction if the ambient field intensity is shifted more than ∼ 25% away from local ambient values (W. Wiltschko, 1972), which would stop them from using this compass over geomagnetic anomalies. From our human experiments to date, we suspect that alpha-ERD occurs in our participants mainly in response to geomagnetic fields that reflect something close to “normal” in the Northern Hemisphere where the North-seeking field vector tilts downwards. This would explain why field rotations with a static upwards component produced little response in Northern Hemisphere participants. Conducting similar experiments on participants born and raised in other geographic regions (such as in the Southern Hemisphere or on the Geomagnetic Equator) could test this hypothesis.

Another question vis-à-vis response selectivity is why downwards-directed CCW (DecDn.CCW.N), but not CW (DecDn.CW.N), rotations elicited alpha-ERD. The bias could arise either at the receptor level or at higher processing levels. The structure and function of the magnetoreceptor cells are unknown, but biological structures exhibit chirality (right-or left-handedness) at many spatial scales – from individual amino acids to folded protein assemblies to multicellular structures. If such mirror asymmetries exist in the macromolecular complex interfacing with magnetite, they could favor the transduction of one stimulus over its opposite. Alternatively, higher-level cognitive processes could tune the neural response towards counterclockwise rotations without any bias at the receptor level. As of this writing, we cannot rule out the possibility that some fraction of humans may have a CW response under this or other experimental paradigms, just as some humans are left-instead of right-handed. We also cannot rule out the existence of a separate neural response to CW rotations that is not reflected in the alpha-ERD signature that we assay here.

The functional significance of the divergent responses to CW and CCW is also unclear. It may simply arise as a byproduct during the evolution and development of other mirror asymmetries (such as north-up vs. north-down), which serve a clearer functional, ecologically relevant purpose with a lower biological cost. It may also be that the alpha-ERD response reflects non-directional information, such as a warning of geomagnetic anomalies, which can expose a navigating animal to sudden shifts of the magnetic field comparable to those used in our experiments. For example, volcanic or igneous terranes are prone to remagnetization by lightning strikes, which produce magnetic fields powerful enough to leave local, 1-10 m scale remnant (permanent) magnetizations strong enough to warp the otherwise uniform local geomagnetic field. A large-scale example is the Vredefort Dome area in South Africa (Carporzen, Weiss, Gilder, Pommier, & Hart, 2012) where lightning remagnetization has been studied extensively. Such anomalies are common in areas with volcanic or igneous basement rock and can be located by simply wandering around with a hand compass held level at waist height and observing abnormal swings of the compass needle from magnetic north. An animal moving through isolated features of this sort would experience paired shifts; the magnetic field direction and intensity would change as the anomaly is entered and then return to normal upon exiting. If the magnetosensory system evolved in the brain as a warning signal against using the magnetic field for long-range navigation while passing through local field anomalies, sensitivity to only one directional excursion is needed. Future experiments could test this speculation by sweeping field intensity through values matching those of lightning-strike and other anomalies to check for asymmetric patterns of alpha desynchronization.

A further question is whether the response asymmetry occurs only in passive experiments when participants experience magnetic stimulation without making use of the information or also in active experiments with a behavioral task, such as judging the direction or rotation of the magnetic field. Behavioral tasks with EEG recording could be used to explore the magnetosensory system in more detail and may uncover the unknown function of the observed response and its asymmetry.

### General Discussion

As noted above, many past attempts have been made to test for the presence of human magnetoreception using *behavioral* assays, but the results were inconclusive. To avoid the cognitive and behavioral artifacts inherent in testing weak or subliminal sensory responses, we decided to use EEG techniques to see directly whether or not the human brain has passive responses to magnetic field changes. Our results indicate that human brains are indeed collecting and selectively processing directional input from magnetic field receptors. These give rise to a brain response that is selective for field direction and rotation with a pattern of neural activity that is measurable at the group level and repeatable in strongly-responding individuals. Such neural activity is a necessary prerequisite for any subsequent behavioral expression of magnetoreception, but such magnetically-triggered neural activity does not demand that the magnetic sense be expressed behaviorally or enter an individual’s conscious awareness.

The fact that alpha-ERD is elicited in a specific and sharply delineated pattern allows us to make inferences regarding the biophysical mechanisms of signal transduction. Notably, the alpha-ERD response differentiated clearly between sets of stimuli differing only by their static or polar components. Electrical induction, electrical artifacts and quantum compass mechanisms are totally insensitive to these components and cannot account for the selectivity of brain responses. Indeed, while birds have evolved a method of navigation that would allow them to navigate by combining a non-polar magnetic sense with gravity, that strategy would not be able to distinguish our test stimuli (see section 8 of the *Extended Materials and Methods).* In contrast, ferromagnetic mechanisms can be highly sensitive to both static and polar field components, and could distinguish our test stimuli with differing responses. Finally, magnetite-based mechanisms for navigation have been characterized in animals through neurophysiological (Walker et al., 1997), histological (Diebel, Proksch, Green, Neilson, & Walker, 2000) and pulse-remagnetization studies (Beason, Wiltschko, & Wiltschko, 1997; Ernst & Lohmann, 2016; R. A. Holland, 2010; R.A. Holland & Helm, 2013; R. A. Holland, Kirschvink, Doak, & Wikelski, 2008; Irwin & Lohmann, 2005; J.L. Kirschvink & Kobayashi-Kirschvink, 1991; Munro, Munro, Phillips, Wiltschko, & Wiltschko, 1997; Munro, Munro, Phillips, & Wiltschko, 1997; W. Wiltschko, Ford, Munro, Winklhofer, & Wiltschko, 2007; W. Wiltschko, Munro, Beason, Ford, & Wiltschko, 1994; W. Wiltschko, Munro, Ford, & Wiltschko, 1998, 2009; W. Wiltschko, Munro, Wiltschko, & Kirschvink, 2002; W. Wiltschko & R. Wiltschko, 1995), and biogenic magnetite has been found in human tissues (Dunn et al., 1995; Gilder et al., 2018; J. L. Kirschvink, Kobayashi-Kirschvink, & Woodford, 1992; Kobayashi & Kirschvink, 1995; Maher et al., 2016; Schultheiss-Grassi, Wessiken, & Dobson, 1999).

These data argue strongly for the presence of geomagnetic transduction in humans, similar to those in numerous migratory and homing animals. Single-domain ferromagnetic particles such as magnetite are directly responsive to both time-varying and static magnetic fields and are sensitive to field polarity. At the cellular level, the magnetomechanical interaction between ferromagnetic particles and the geomagnetic field is well above thermal noise (J.L. Kirschvink & Gould, 1981; J. L. Kirschvink, Winklhofer, & Walker, 2010), stronger by several orders of magnitude in some cases (Eder et al., 2012). In many animals, magnetite-based transduction mechanisms have been found and shown to be necessary for navigational behaviors, through neurophysiological and histological studies (Diebel et al., 2000; Walker et al., 1997). A natural extension of this study would be to apply the pulse-remagnetization methods used in animals to directly test for a ferromagnetic transduction element in humans. In these experiments, a brief magnetic pulse causes the magnetic polarity of the single-domain magnetite crystals to flip. Following this treatment, the physiological and behavioral responses to the geomagnetic field are expected to switch polarity. These experiments could provide measurements of the microscopic coercivity of the magnetite crystals involved and hence make predictions about the physical size and shape of the crystals involved (Diaz Ricci & Kirschvink, 1992 203), and perhaps their physiological location.

Previous attempts to detect human magnetoreception may have been confounded by a number of factors. Response specificity and neural tuning to the local environment (Block, 1992) make it likely that tests using stimuli outside the environmental range would likely fail, and past computational methods were not as good at isolating the neural activity studied here (Boorman et al., 1999; Sastre et al., 2002), further discussed in section 9 of the *Extended Materials and Methods,* below. Other experiments were conducted in unshielded conditions and may have been subject to radio-frequency noise which has been shown to shut down magnetoreceptivity in birds and other animals (Engels et al., 2014; Landler, Painter, Youmans, Hopkins, & Phillips, 2015; Tomanova & Vacha, 2016; R. Wiltschko et al., 2015).

At this point, our observed reduction in alpha-band power is a clear neural signature for cortical processing of the geomagnetic stimulus, but its functional significance is unknown. In form, the activity is an alpha-ERD response resembling those found in other EEG investigations of sensory and cognitive processing. However, the alpha-ERD responses found in literature take on a range of different spatiotemporal forms and are associated with a variety of functions. It is likely that the alpha-ERD seen here reflects the sudden recruitment of neural processing resources, as this is a finding common across studies. But more research will be needed to see if and how it relates more specifically to previously studied processes such as memory access or recruitment of attentional resources.

Further, alpha-ERD probably represents only the most obvious signature of neural processing arising from geomagnetic input. A host of upstream and downstream processes need to be investigated to reveal the network of responses and the information they encode. Responses independent from the alpha-ERD signature will likely emerge, and those might show different selectivity patterns and reflect stimulus features not revealed in this study. Does human magnetoreceptive processing reflect a full representation of navigational space? Does it contain certain warning signals regarding magnetic abnormalities? Or have some aspects degenerated from the ancestral system? For now, alpha-ERD remains a blank signature for a wider, unexplored range of magnetoreceptive processing.

Future experiments should examine how magnetoreceptive processing interacts with other sensory modalities in order to determine field orientation. Our experimental results suggest the combination of a magnetic and a positional cue (e.g. reacting differently to North-up and North-down fields). However, we cannot tell if this positional cue uses a reference frame set by gravity sensation or is aligned with respect to the human body. In birds, orientation behavior reflects a magnetic inclination compass that identifies the steepest angle of magnetic field dip with respect to gravity (R. Wiltschko & W. Wiltschko, 1995; W. Wiltschko, 1972), and this compass can operate at dips as shallow as 5° from horizontal (Schwarze et al., 2016). Because magnetism and gravity are distinct, non-interacting forces of nature, the observed behavior must arise from processing of neural information from separate sensory systems (J. L. Kirschvink et al., 2010). Evolution has driven many of the known sensory systems down to their physical detection limits with astounding specificity (Block, 1992). Gravitational information is known to arise in the utricle and saccule of the vertebrate vestibular system due to the motions of dense biominerals activating hair cells (Lopez & Blanke, 2011), and a magnetite-based magnetosensory organ has been localized at the cellular level in fish (Diebel et al., 2000; Eder et al., 2012; Walker et al., 1997). The neural processing of magnetic with gravitational sensory cues could perhaps be addressed by modifying the test chamber to allow the participant to rest in different orientations with respect to gravity or by running the experiment in the zero-gravity environment of the international space station.

In the participant pool, we found several highly responsive individuals whose alpha-ERD proved to be stable across time: 4 participants responded strongly at the p<0.01 level in repeated testing over weeks or months. Repeatability in individual participants suggests that the alpha-ERD did not arise due to chance fluctuations in a single run, but instead reflects a consistent individual characteristic, measurable across multiple runs. A wider survey of individuals could reveal genetic/developmental or other systematic differences underlying these individual differences.

The range of individual responses may be partially attributed to variation in basic alpha-ERD mechanisms, rather than to underlying magnetoreceptive processing. However, some participants with high resting alpha power showed very little alpha-ERD to the magnetic field rotations, suggesting that the extent of magnetoreceptive processing itself varies across individuals. If so, distinct human populations may be good targets for future investigation. For example, studies of comparative linguistics have identified a surprising number of human languages that rely on a cardinal system of environmental reference cues (e.g. North, South, East, West) and lack egocentric terms like front, back, left, and right (Haviland, 1998; Levinson, 2003; Meakins, 2011; Meakins & Algy, 2016; Meakins, Jones, & Algy, 2016). Native speakers of such languages would (e.g.) refer to a nearby tree as being to their North rather than being in front of them; they would refer to their own body parts in the same way. Individuals who have been raised from an early age within a linguistic, social and spatial framework using cardinal reference cues might have made associative links with geomagnetic sensory cues to aid in daily life; indeed, linguists have suggested a human magnetic compass might be involved (Levinson, 2003). It would be interesting to test such individuals using our newly-developed methods to see if such geomagnetic cues might already be within their conscious awareness, aiding their use of the cardinal reference system. In turn, such experiments might guide the development of training procedures to enhance geomagnetic sensitivity in individuals raised in other language and cultural groups, advancing more rapidly studies on the nature of a human magnetic sense.

In the 198 years since Danish physicist Hans Christian Ørsted discovered electromagnetism (March 1820), human technology has made ever-increasing use of it. Most humans no longer need to rely on an internal navigational sense for survival. To the extent that we employ a sense of absolute heading in our daily lives, external cues such as landmarks and street grids can provide guidance. Even if an individual possesses an implicit magnetoreceptive response, it is likely to be confounded by disuse and interference from our modern environment. A particularly pointed example is the use of strong permanent magnets in both consumer and aviation headsets, most of which produce static fields through the head several times stronger than the ambient geomagnetic field. If there is a functional significance to the magnetoreceptive response, it would have the most influence in situations where other cues are impoverished, such as marine and aerial navigation, where spatial disorientation is a surprisingly persistent event (Poisson & Miller, 2014). The current alpha-ERD evidence provides a starting point to explore functional aspects of magnetoreception, by employing various behavioral tasks in variety of sensory settings.

## Conclusion

We conclude that at least some modern humans transduce changes in Earth-strength magnetic fields into an active neural response. Given the known presence of highly-evolved geomagnetic navigation systems in species across the animal kingdom, it is perhaps not surprising that we might retain at least some functioning neural components, especially given the nomadic hunter/gatherer lifestyle of our not-too-distant ancestors. The full extent of this inheritance remains to be discovered.

### Extended Materials and Methods

Detailed additional instructions concerning the custom-built equipment and instrumentation are provided below. All experiments were performed in accordance with relevant guidelines and regulations following NIH protocols for human experimentation, as reviewed and approved periodically by the Caltech Administrative Committee for the Protection of Human Subjects (Caltech IRB, protocols 13-0420, 17-0706, and 17-0734). All methods were carried out in accordance with relevant guidelines and regulations. Informed consent using forms approved by the Institutional Review Board was obtained from all subjects. No subjects under the age of 18 were used in these experiments.

#### 1. Magnetic Exposure Facility

We constructed a six-sided Faraday cage shown in Figs. 1 and 5 out of aluminum, chosen because of: (1) its high electrical conductivity, (2) low cost and (3) lack of ferromagnetism. The basic structure of the cage is a rectangular 2.44 m × 2.44 m × 2.03 m frame made of aluminum rods, 1.3 cm by 1.3 cm square in cross-section, shown in Fig. 5A. Each of the cage surfaces (walls, floor and ceiling) have four rods (two vertical and two horizontal) bounding the perimeter of each sheet. On the cage walls three vertical rods are spaced equally along the inside back of each surface, and on the floor and ceiling three horizontal rods are similarly spaced, forming an inwards-facing support frame. This frame provides a conductive chassis on which overlapping, 1 mm thick aluminum sheets (2.44 m long and 0.91 m wide) were attached using self-threading aluminum screws at ∼0.60 m intervals with large overlaps between each sheet. In addition, we sealed the seams between separate aluminum panels with conductive aluminum tape. The access door for the cage is a sheet of aluminum that is fastened with a 2.4 m long aluminum hinge on the East-facing wall such that it can swing away from the cage and provide an entrance/exit. Aluminum wool has been affixed around the perimeter of this entrance flap to provide a conductive seal when the flap is lowered (e.g. the cage is closed). Ventilation is provided via a ∼3 m long, 15 cm diameter flexible aluminum tube (Fig. 5E) that enters an upper corner of the room and is connected to a variable-speed ceiling-mounted fan set for a comfortable but quiet level of airflow. The end of the tube in contact with the Faraday cage is packed loosely with aluminum wool that allows air to pass and provides electrical screening. LED light strips (Fig. 5H) provide illumination for entrance and exit. These lights are powered by a contained lithium ion battery housed in an aluminum container attached at the top end of the Faraday cage, adjacent to the entrance of the ventilation air duct (seen as the red battery in Fig. 5E).

**Fig 5.**
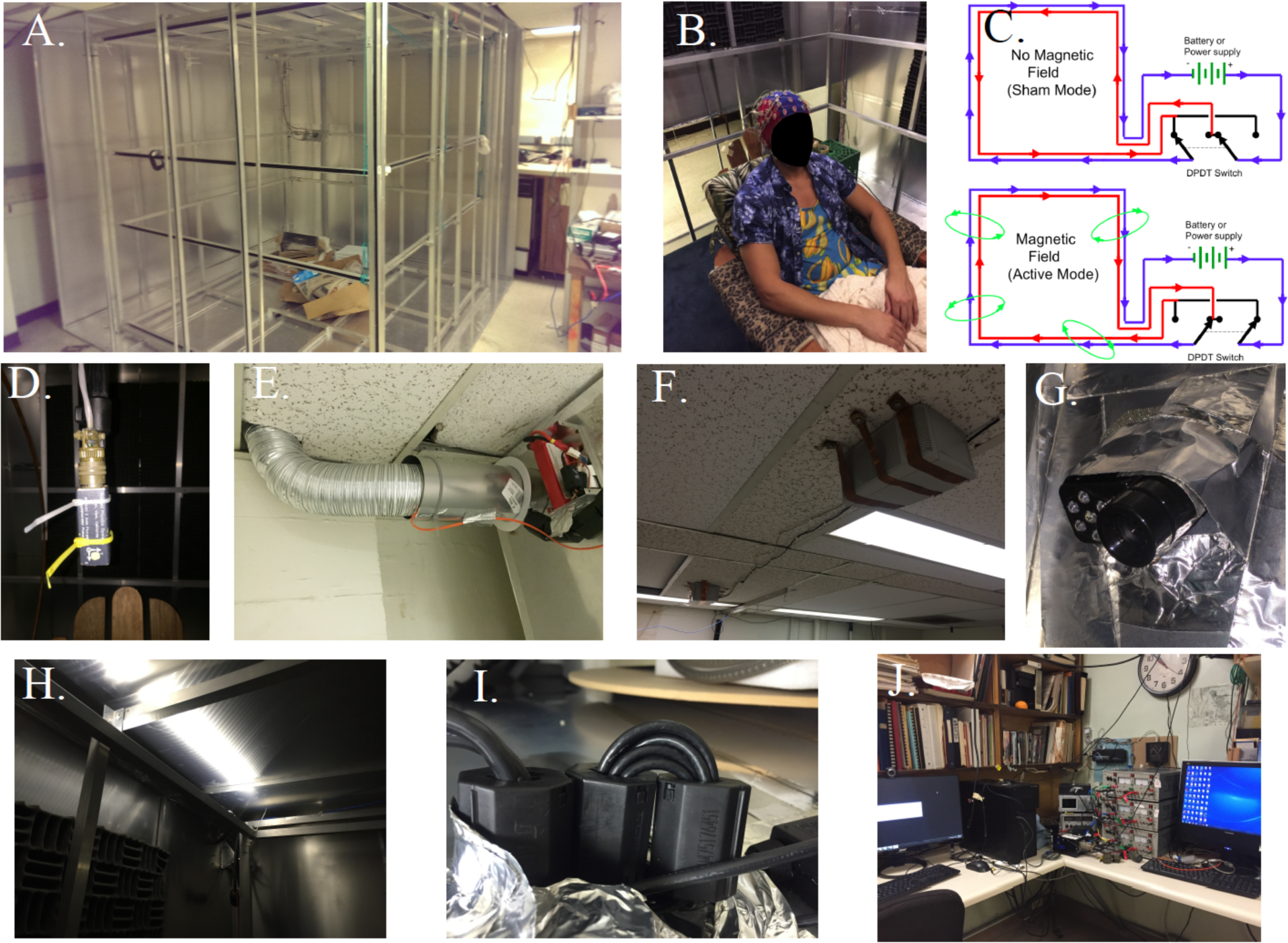
Additional images of critical aspects of the human magnetic exposure facility at Caltech. A. Partially complete assembly of the Faraday cage (summer of 2014) showing the nested set of orthogonal, Merritt square four-coils (Merritt et al., 1983) with all but two aluminum walls of the Faraday cage complete. B. Image of a participant in the facility seated in a comfortable, non-magnetic wooden chair and wearing the 64-lead BioSim^™^ EEG head cap. The EEG sensor leads are carefully braided together to minimize electrical artifacts. The chair is on a raised wooden platform that is isolated mechanically from the magnet coils and covered with a layer of synthetic carpeting; the height is such that the participant’s head is in the central area of highest magnetic field uniformity. C. Schematic of the double-wrapped control circuits that allow active-sham experiments (J.L Kirschvink, 1992). In each axis of the coils, the four square frames are wrapped in series with two discrete strands of insulated copper magnet wire and with the number of turns and coil spacing chosen to produce a high-volume, uniform applied magnetic field (Merritt et al., 1983). Reversing the current flow in one of the wire strands via a double-pole-double-throw (DPDT) switch results in cancellation of the external field with virtually all other parameters being the same. This scheme is implemented on all three independently controlled coil axes (Up/Down, East/West and North/South). D. Fluxgate magnetometer (Applied Physics Systems 520A) three-axis magnetic field sensor attached to a collapsing carbon-fiber camera stand mount. At the start of each session the fluxgate is lowered to the center of the chamber for an initial current / control calibration of the ambient geomagnetic field. It is then raised to a position about 30 cm above the participant’s head during the following experimental trials, and the three-axis magnetic field readings are recorded continuously in the same fashion as the EEG voltage signals. E. Air duct. A 15 cm diameter aluminum air duct ∼2 meters long connects a variable-speed (100 W) electric fan to the upper SE corner of the experimental chamber; this is also the conduit used for the major electrical cables (power for the magnetic coils, sensor leads for the fluxgate, etc.). F. & G. An intercom / video monitoring system was devised by mounting a computer-controlled loudspeaker (F) outside the Faraday shield on the ceiling North of the chamber coupled with (G) a USB-linked IR video camera / microphone system mounted just inside the shield. Note the conductive aluminum tape shielding around the camera to reduce Rf interference. During all experimental trials a small DPDT relay located in the control room disconnects the speaker from computer and directly shorts the speaker connections. A second microphone in the control room can be switched on to communicate with the participant in the experimental chamber, as needed. An experimenter monitors the audio and video of participants at all times, as per Caltech IRB safety requirements. H. LED lights, 12 VDC array, arranged to illuminate from the top surface of the magnetic coils near the ceiling of the chamber. These are powered by rechargeable 11.1 V lithium battery packs (visible in E) and controlled by an external switch. I. Ferrite chokes. Whenever possible, these are mounted in a multiple-turn figure-eight fashion (Counselman, 2013) on all conductive wires and cables entering the shielded area and supplemented with grounded aluminum wool when needed. J. Image of the remote control area including (from left to right): the PC for controlling the coils, the DPDT switches for changing between active and sham modes, the fluxgate control unit, the three power amplifiers that control the current in the remote coil room, and the separate PC that records the EEG data. Participants seated in the experimental chamber do not report being able to hear sounds from the control room and *vice versa.*

In all experiment sessions, power to the lights was switched off. A small USB-powered infrared camera and microphone assembly (Fig. 5G) mounted just inside the cage on the North wall allows audiovisual monitoring of participants inside the room. Instructions to the participants are given from a pair of speakers mounted outside the Faraday cage (Fig. 5F), controlled remotely by experimenters and electrically shorted by a computer-controlled TTL relay when not in use. Acoustic foam panels are attached to the vertical walls to dampen echoes within the chamber as well as to reduce the amplitude of external sound entering the chamber. To complete the Faraday shielding, we grounded the cage permanently at one corner with a 2.6 mm diameter (10 AWG) copper wire connected to the copper plumbing in the sub-basement of the building. RMS noise measurements from the cage interior using a Schwarzbeck Mess^™^ Elektronik FMZB 1513 B-component active loop Rf antenna, a RIGOL^™^ DSA815/E-TG spectrum analyzer, and a Tektronix^™^ RSA503A signal analyzer indicated residual noise interference below 0.01 nT, in the frequency range from 9 kHz to 10 MHz.

Electrical cables entering the Faraday cage pass through a side gap in the aluminum ventilation duct and then through the aluminum wool. Rf interference is blocked further on all electrical cables entering the room using pairs of clip-on ferrite chokes (Fair-Rite^™^ material #75, composed of MnZn ferrite, designed for low-frequency EMI suppression, referred from here-on as ferrite chokes) and configured where possible using the paired, multiple-loop “pretty-good choke” configuration described by Counselman (Counselman, 2013) (Fig. 5I). Inside the shielded space are located a three-axis set of square coils approximately 2 m on edge following the Merritt *et al.* four-coil design (Merritt et al., 1983) (using the 59/25/25/59 coil winding ratio) that provides remarkably good spatial uniformity in the applied magnetic field (12 coils total, four each in the North/South, East/West, and Up/Down orientations as seen in Fig. 5A). The coils are double-wrapped inside grounded aluminum U-channels following a design protocol that allows for full active-field and sham exposures (J.L Kirschvink, 1992); they were constructed by Magnetic Measurements, Ltd., of Lancashire, U.K. (http://www.magnetic-measurements.com). This double-wrapped design gives a total coil winding count of 118/50/50/118 for all three-axes coil sets.

To provide a working floor isolated from direct contact with the coils, we suspended a layer of ∼2 cm thick plywood sheets on a grid work of ∼ 10 × 10 cm thick wooden beams that rested on the basal aluminum plate of the Faraday shield that are held together with brass screws. We covered this with a layer of polyester carpeting on top of which we placed a wooden platform chair for the participants (Fig. 5B). Non-magnetic bolts and screws were used to fasten the chair together, and a padded foam cushion was added for comfort. The chair is situated such that the head and upper torso of most participants fit well within the ∼1 m^3^ volume of highly uniform magnetic fields produced by the coil system (J.L Kirschvink, 1992) while keeping the participants a comfortable distance away from direct contact with the Merritt coils.

We suspended the three-axis probe of a fluxgate magnetometer (Applied Physics Systems^™^ model 520A) on a non-magnetic, carbon-fiber, telescoping camera rod suspended from the ceiling of the Faraday cage (Fig. 5D). This was lowered into the center of the coil system for initial calibration of field settings prior to experiments and then raised to the edge of the uniform field region to provide continuous recording of the magnetic field during experiments. Power cables for the coils and a data cable for the fluxgate sensor pass out of the Faraday cage through the ventilation shaft, through a series of large Rf chokes (Counselman, 2013), a ceiling utility chase in the adjacent hallway, along the wall of the control room, and finally down to the control hardware. The control hardware and computer are located ∼20 m away from the Faraday cage through two heavy wooden doors and across a hallway that serve as effective sound dampeners such that participants are unable to directly hear the experimenters or control equipment and the experimenters are unable to directly hear the participant.

In the remote-control room, three bipolar power amplifiers (Kepco^™^ model BOP-100-1MD) control the electric power to the coil systems (Fig. 5J) and operate in a mode where the output current is regulated proportional to the control voltage, thereby avoiding a drop in current (and magnetic field) should the coil resistance increase due to heating. Voltage levels for these are generated using a 10k samples per channel per second, 16-bit resolution, USB-controlled, analog output DAQ device (Measurement Computing^™^ Model USB-3101FS), controlled by the desktop PC. This same PC controls the DC power supply output levels, monitors and records the Cartesian orthogonal components from the fluxgate magnetometer, displays video of the participant (recordings of which are not preserved per IRB requirements), and is activated or shorted, via TTL lines, to the microphone/speaker communication system from the control room to the experimental chamber. As the experimenters cannot directly hear the participant and the participant cannot directly hear the experimenters, the microphone and speaker system are required (as per Caltech Institute Review Board guidelines) to ensure the safety and comfort of the participant as well as to pass instructions to the participant and answer participants’ questions before the start of a block of experiments. The three-axis magnet coil system can produce a magnetic vector of up to 100 μT intensity (roughly 2-3X the background strength in the lab) in any desired direction with a characteristic RL relaxation constant of 79-84 ms (inductance and resistance of the four coils in each axis vary slightly depending on the different coil-diameters for each of the three nested, double-wrapped coil-set axes). Active/Sham mode was selected prior to each run via a set of double-pole-double-throw (DPDT) switches located near the DC power supplies. These DPDT switches are configured to swap the current direction flowing in one strand of the bifilar wire with respect to the other strand in each of the coil sets (J.L Kirschvink, 1992) (Fig. 5C). Fluxgate magnetometer analog voltage levels were digitized and streamed to file via either a Measurement Computing^™^ USB 1608GX 8-channel (differential mode) analog input DAQ device, or a Measurement Computing^™^ USB 1616HS-2 multifunction A/D, D/A, DIO DAQ device connected to the controller desktop PC. Fluxgate analog voltage signal levels were sampled at 1024 or 512 Hz. Although the experimenter monitors the audio/video webcam stream of the participants continuously, as per Caltech IRB safety requirements, while they are in the shielded room the control software disconnects the external speakers (in the room that houses the experimental Faraday cage and coils) and shorts them to electrical ground during all runs to prevent extraneous auditory cues from reaching the participants. Light levels within the experimental chamber during experimental runs were measured using a Konica-Minolta CS-100A luminance meter, which gave readings of zero (below 0.01 ± 2% cd/m^2^).

#### 2. Participants

Participants were 34 adult volunteers (24 male, 12 female) recruited from the Caltech population. This participant pool included persons of European, Asian, African and Native American descent. Ages ranged from 18 to 68 years. Each participant gave written informed consent of study procedures approved by the Caltech Institutional Review Board (Protocols 13-0420, 17-0706, and 17-0734).

#### 3. Experimental Protocol

In the experiment, participants sat upright in the chair with their eyes closed and faced North (defined as 0° declination in our magnetic field coordinate reference frame). The experimental chamber was dark, quiet and isolated from the control room during runs. Each run was ∼7 minutes long with up to eight runs in a ∼1 hour session. The magnetic field was rotated over 100 milliseconds every 2-3 seconds, with constant 2 or 3 s inter-trial intervals in early experiments and pseudo-randomly varying 2-3 s intervals in later experiments. Participants were blind to Active vs. Sham mode, trial sequence and trial timing. During sessions, auditory tones signaled the beginning and end of experiments and experimenters only communicated with participants once or twice per session to update the number of runs remaining. When time allowed, Sham runs were matched to Active runs using the same software settings. Sham runs are identical to Active runs but are executed with the current direction switches set to anti-parallel. This resulted in no observable magnetic field changes throughout the duration of a Sham run with the local, uniform, static field produced by the double-wrapped coil system in cancellation mode (J.L Kirschvink, 1992).

Two types of trial sequences were used: (1) a 127-trial Gold Sequence with 63 FIXED trials and 64 SWEEP trials evenly split between two rotations (32 each), and (2) various 150-trial pseudorandom sequences with 50 trials of each rotation interspersed with 50 FIXED trials to balance the number of trials in each of three conditions. All magnetic field parameters were held constant during FIXED trials, while magnetic field *intensity* was held constant during inclination or declination rotations. In inclination experiments (Fig. 2A of the main text), the vertical component of the magnetic field was rotated upwards and downwards between ±55°, ±60°, or ±75° (Inc.UP and Inc.DN, respectively); data collected from runs with each of these inclination values were collapsed into a single set representative of inclination rotations between steep angles. In each case, the horizontal component was steady at 0° declination (North; Inc.UP.N and Inc.DN.N). Two types of declination experiments were conducted, designed to test the quantum compass and electrical induction hypotheses. As the quantum compass can only determine the axis of the field and not polarity, we compared a pair of declination experiments in which the rotating vectors were swept down to the North (DecDn.N) and up to the South (DecUp.S), providing two symmetrical antiparallel data sets (Fig. 2B of the main text). In the DecDn.N experiments, the vertical component was held constant and downwards at +60° or +75°, while the horizontal component was rotated between NE (45°) and NW (315°), along a Northerly arc (DecDn.CW.N and DecDn.CCW.N). In DecUp.S experiments, the vertical component was held upwards at −60° or −75°, while the horizontal component was rotated between SW (225°) and SE (135°) along a Southerly arc (DecUp.CW.S and DecUp.CCW.S). Again, runs with differing inclination values were grouped together as datasets with steep downwards or steep upwards inclination. To test the induction hypothesis, we paired the DecDn.N sweeps with a similar set, DecUp.N, as shown on Fig. 2C. These two conditions only differ in the direction of the vertical field component; rotations were between NE and NW in both experiments (DecDn.CW.N, DecDn.CCW.N, DecUp.CW.N and DecUp.CCW.N). Hence, any significant difference in the magnetosensory response eliminates induction as a mechanism.

#### 4. EEG Recording

EEG was recorded using a BioSemi^™^ ActiveTwo system with 64 electrodes following the International 10-20 System (Nuwer et al., 1998). Signals were sampled at 512 Hz with respect to CMS/DRL reference at low impedance <1 ohm and bandpass-filtered from 0.16-100 Hz. To reduce electrical artifacts induced by the time-varying magnetic field, EEG cables were bundled and twisted 5 times before plugging into a battery-powered BioSemi^™^ analog/digital conversion box. Digitized signals were transmitted over a 30 m, non-conductive, optical fiber cable to a BioSemi^™^ USB2 box located in the control room ∼20 m away where a desktop PC (separate from the experiment control system) acquired continuous EEG data using commercial ActiView^™^ software. EEG triggers signaling the onset of magnetic stimulation were inserted by the experiment control system by connecting a voltage timing signal (0 to 5 V) from its USB-3101FS analog output DAQ device. The timing signal was sent both to the Measurement Computing USB-1608GX (or USB-1616HS-2) analog input DAQ device, used to sample the magnetic field on the experiment control PC and a spare DIO voltage input channel on the EEG system’s USB2 DAQ input box, which synchronized the EEG data from the optical cable with the triggers cued by the controlling desktop PC. This provided: (1) a precise timestamp in continuous EEG whenever electric currents were altered (or in the case of FIXED trials, when the electric currents could have been altered to sweep the magnetic field direction, but were instead held constant) in the experimental chamber, and (2) a precise correlation (±2 ms, precision determined by the 512 samples per second digital input rate of the BioSemi^™^ USB2 box) between fluxgate and EEG data.

#### 5. EEG Analysis

Raw EEG data were extracted using EEGLAB^™^ toolbox for MATLAB^™^ and analyzed using custom MATLAB^™^ scripts. Trials were defined as 2- or 3-s epochs from −0.75 s pre-stimulus to +1.25 or +2.25 s post-stimulus, with a baseline interval from −0.5 s to −0.25 s pre-stimulus. Time/frequency decomposition was performed for each trial using Fast Fourier Transform (MATLAB^™^ function *fft*) and Morlet wavelet convolution on 100 linearly-spaced frequencies between 1 and 100 Hz. Average power in an extended alpha band of 6-14 Hz was computed for the pre-stimulus and post-stimulus intervals of all trials, and a threshold of 1.5X the interquartile range was applied to identify trials with extreme values of log alpha power. These trials were excluded from further analysis but retained in the data. After automated trial rejection, event-related potentials (ERPs) were computed for each condition and then subtracted from each trial of that condition to reduce the electrical induction artifact that appeared only during the 100 ms magnetic stimulation interval. This is an established procedure to remove phase-locked components such as sensory-evoked potentials from an EEG signal for subsequent analysis of non-phase-locked, time/frequency power representations. Non-phase-locked power was computed at midline frontal electrode Fz for each trial and then averaged and baseline-normalized for each condition to generate a time/frequency map from −0.25 s pre-stimulus to +1 s or +2 s post-stimulus and 1-100 Hz. To provide an estimate of overall alpha power for each participant, power spectral density was computed using Welch’s method (MATLAB^™^ function *pwelch*) at 0.5 Hz frequency resolution (Welch, 1967).

From individual datasets, we extracted post-stimulus alpha power to test for statistically significant differences amongst conditions at the group level. Because alpha oscillations vary substantially across individuals in amplitude, frequency and stimulus-induced changes, an invariant time/frequency window would not capture stimulus-induced power changes in many participants. In our dataset, individual alpha oscillations ranged in frequency (8 to 12 Hz peak frequency), and individual alpha-ERD responses started around +0.25 to +0.75 s post-stimulus. Thus, we quantified post-stimulus alpha power within an automatically-adjusted time/frequency window for each dataset. First, non-phase-locked alpha power between 6-14 Hz was averaged over all trials regardless of condition. Then, the most negative time/frequency point was automatically selected from the post-stimulus interval between 0 s and +1 or +2 s in this cross-conditional average. The selected point represented the maximum alpha-ERD in the average over all trials with no bias for any condition. A time/frequency window of 0.25 s and 5 Hz was centered (as nearly as possible given the limits of the search range) over this point to define an individualized timing and frequency of alpha-ERD for each dataset. Within the window, non-phase-locked alpha power was averaged across trials and baseline-normalized for each condition, generating a value of alpha-ERD for each condition to be compared in statistical testing.

In early experiments, trial sequences were balanced with nearly equal numbers of FIXED (63) and SWEEP (64) trials, with an equal number of trials for each rotation (e.g. 32 Inc.DN and 32 Inc.UP trials). Later, trial sequences were designed to balance the number of FIXED trials with the number of trials of each rotation (e.g. 50 DecDn.FIXED, 50 DecDn.CCW, and 50 DecDn.CW trials). Alpha-ERD was computed over similar numbers of trials for each condition. For example, when comparing alpha-ERD in the FIXED vs. CCW vs. CW conditions of a declination experiment with 63 FIXED (32 CCW and 32 CW trials) 100 samplings of 32 trials were drawn from the pool of FIXED trials, alpha-ERD was averaged over the subset of trials in each sampling, and the average over all samplings was taken as the alpha-ERD of the FIXED condition. When comparing FIXED vs. SWEEP conditions of an inclination experiment with 50 FIXED, 50 DN, and 50 UP trials, 200 samplings of 25 trials were drawn from each of the DN and UP conditions and the average alpha-ERD over all samplings taken as the alpha-ERD of the SWEEP condition. Using this method, differences in experimental design were reduced, allowing statistical comparison of similar numbers of trials in each condition. The alpha-ERD values for each participant in each condition are shown as histograms for the DecDn (Fig. 6), DecUp (Fig. 7), and Sham declination (Fig. 8) experiments. These values were used in statistical testing at the group level.

**Fig 6.**
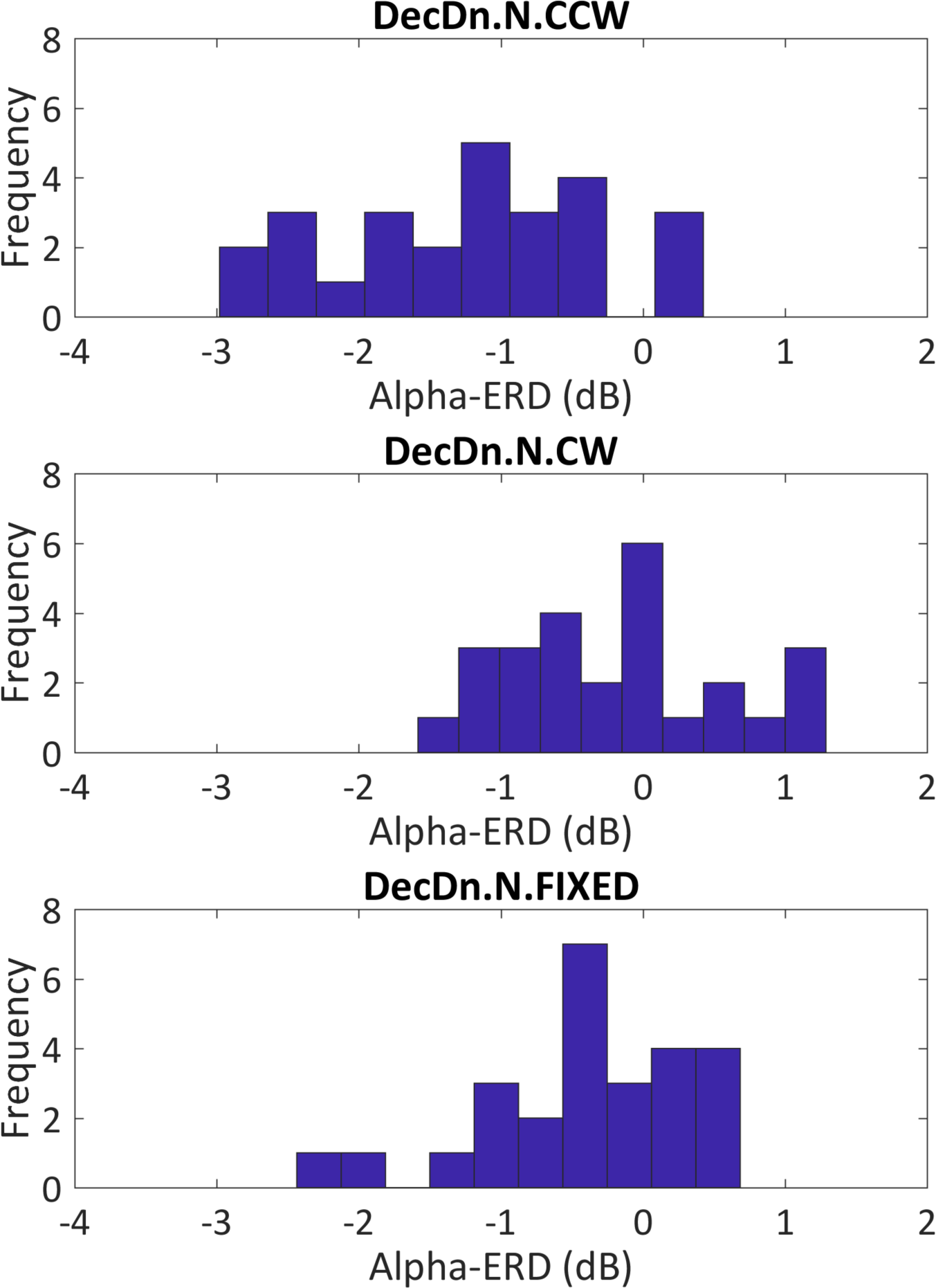
Histogram of alpha-ERD responses over all participants (N=26) in the DecDn experiment. The panels show the histogram of individual responses for each condition. Frequency is given in number of participants. Because we looked for a drop in alpha power following magnetic stimulation, the histograms are shifted towards negative values in all conditions. The CCW condition shows the most negative average in a continuous distribution of participant responses, with the most participants having a >2 dB response.

**Fig 7.**
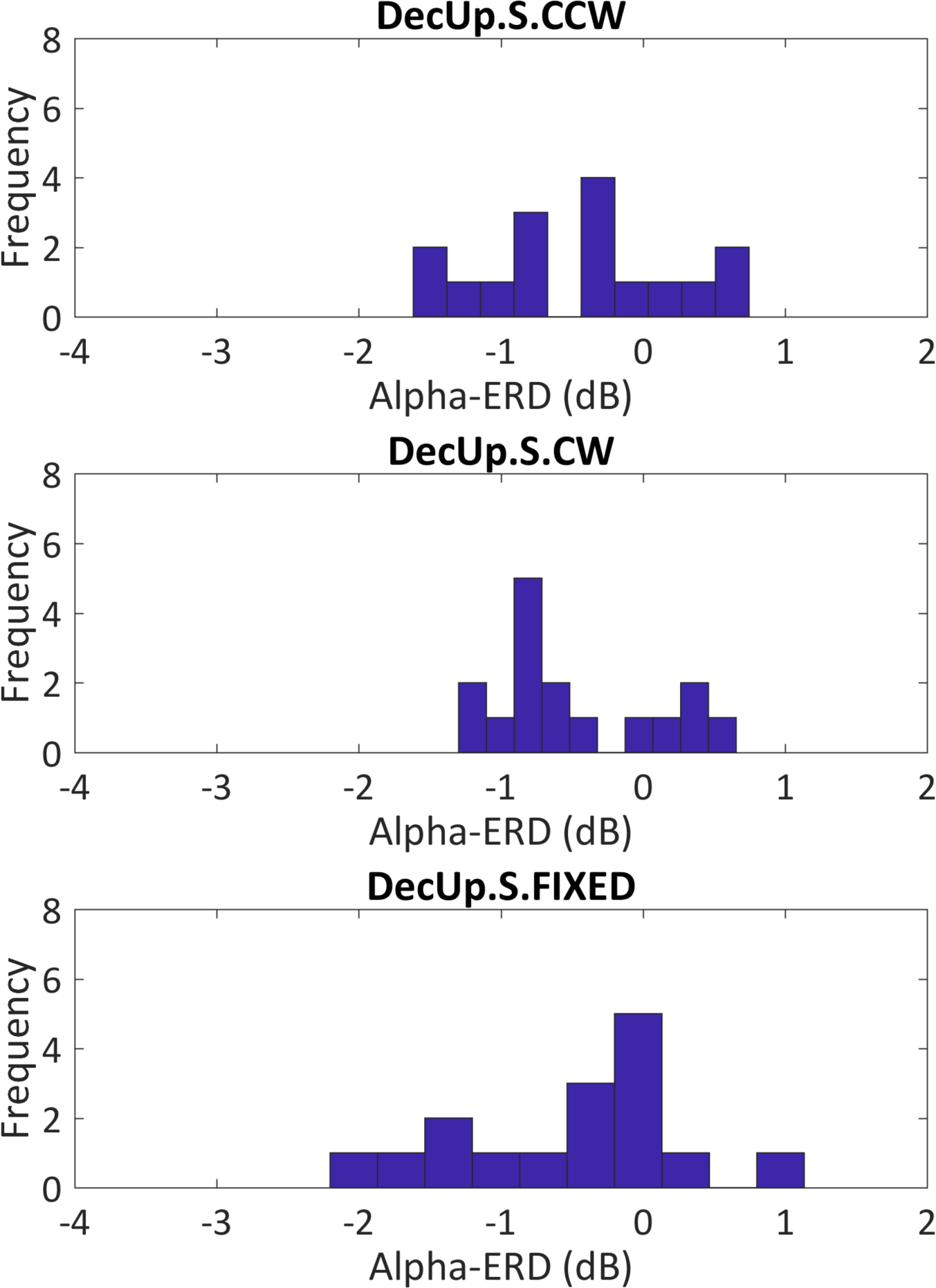
Histogram of alpha-ERD responses over all participants (N=16) in the DecUp experiment. The panels show the histogram of individual responses for each condition. No significant magnetosensory response was observed in any condition, and no clear difference is apparent between the three distributions.

**Fig 8.**
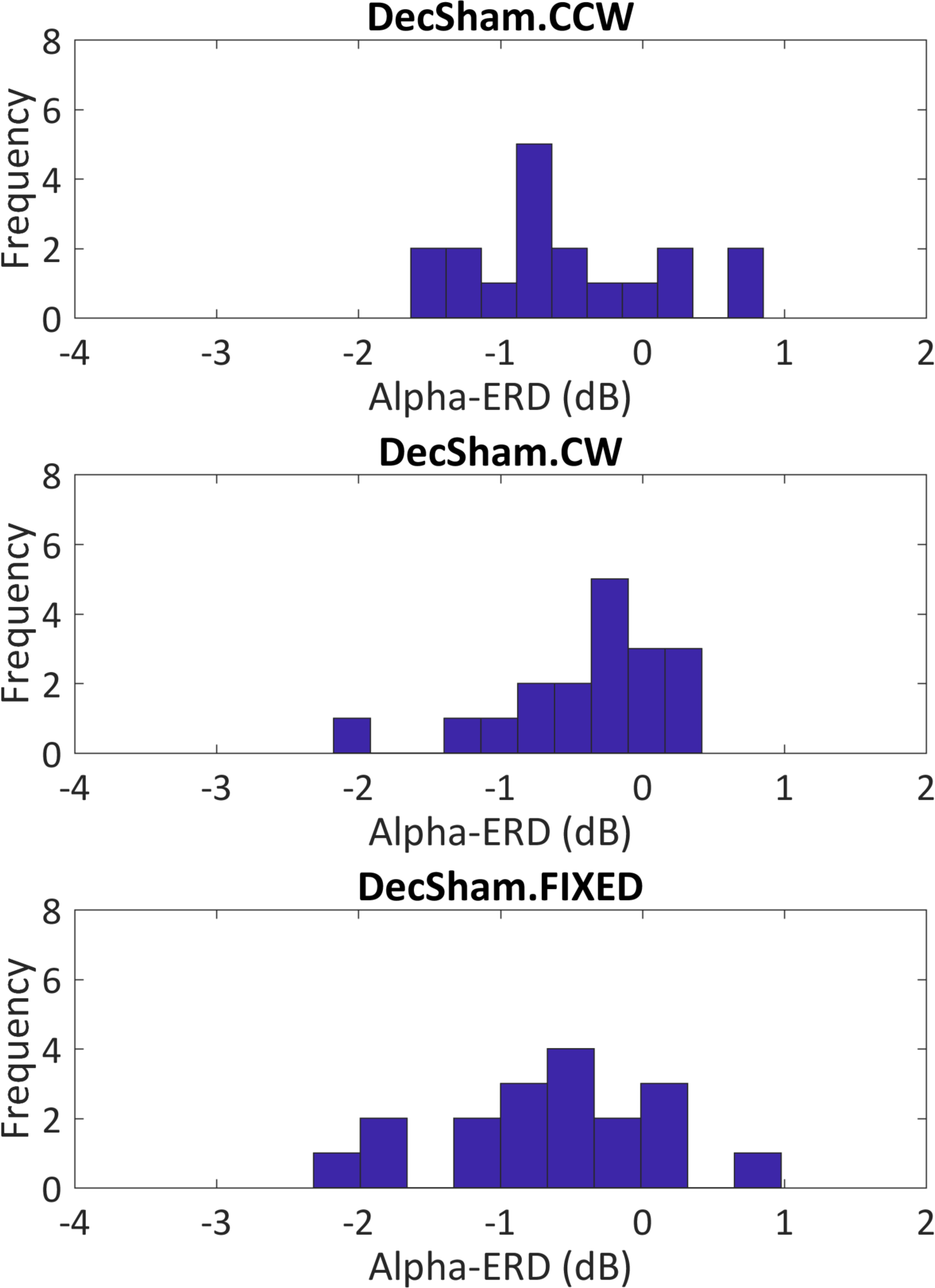
Histogram of alpha-ERD responses over all participants (N=18) in the Sham Declination experiment. The panels show the histogram of individual responses for each condition. No significant magnetosensory response was observed in any condition, and no clear difference is apparent between the three distributions.

Three statistical tests were performed using average alpha-ERD: (1) Inc ANOVA (N=29), (2) DecDn ANOVA (N=26), (3) DecDn/DecUp ANOVA (N=16). For the inclination experiment, data were collected in Active and Sham modes for 29 of 34 participants. Due to time limitations within EEG sessions, sham data could not be collected for every participant, so those participants without inclination sham data were excluded. A two-way repeated-measures ANOVA tested for the effects of inclination rotation (SWEEP vs. FIXED) and magnetic stimulation (Active vs. Sham) on alpha-ERD. Post-hoc testing using the Tukey-Kramer method compared four conditions (Active-SWEEP, Active-FIXED, Sham-SWEEP and Sham-FIXED) for significant differences (Tukey, 1949).

For the DecDn experiment, data were collected from 26 participants in Active mode. A one-way repeated-measures ANOVA tested for the effect of declination rotation (DecDn.CCW vs. DecDn.CW vs. DecDn.FIXED) with post-hoc testing to compare these three conditions. For a subset of participants (N=16 of 26), data was collected from both DecDn and DecUp experiments. The DecUp experiments were introduced in a later group to evaluate the quantum compass mechanism of magnetosensory transduction, as well as in a strongly-responding individual to test the less probable induction hypothesis, as shown in Video 1. For tests of the quantum compass hypothesis, we used the DecDn/DecUp dataset. A two-way repeated-measures ANOVA tested for the effects of declination rotation (DecDn.CCW.N vs. DecDn.CW.N vs. DecUp.CCW.S vs. DecUp.CW.S vs. DecDn.FIXED.N vs. DecUp.FIXED.S) and inclination direction (Inc.DN.N vs Inc.UP.S) on alpha-ERD; data from another strongly-responding individual is shown in Video 2. Post-hoc testing compared six conditions (DecDn.CCW.N, DecDn.CW.N, DecDn.FIXED.N, DecUp.CCW.S, DecUp.CW.S and DecUp.FIXED.S).

Within each group, certain participants responded strongly with large alpha-ERD while others lacked any response to the same rotations. To establish whether a response was consistent and repeatable, we tested individual datasets for significant post-stimulus power changes in time/frequency maps between 0 to +2 or +3 s post-stimulus and 1-100 Hz. For each dataset, 1000 permutations of condition labels over trials created a null distribution of post-stimulus power changes at each time/frequency point. The original time/frequency maps were compared with the null distributions to compute a p-value at each point. False discovery rate correction for multiple comparisons was applied to highlight significant post-stimulus power changes at the p<0.05 and p<0.01 statistical thresholds (Benjamini & Hochberg, 1995). Fig. 9 shows repeated runs (Run #1 and Run #2) of two different participants (A and B) in the DecDn experiment. The outlined clusters indicate significant power changes following magnetic field rotation. In each case, the significant clusters are similar in timing and bandwidth across runs up to six months apart.

**Fig 9.**
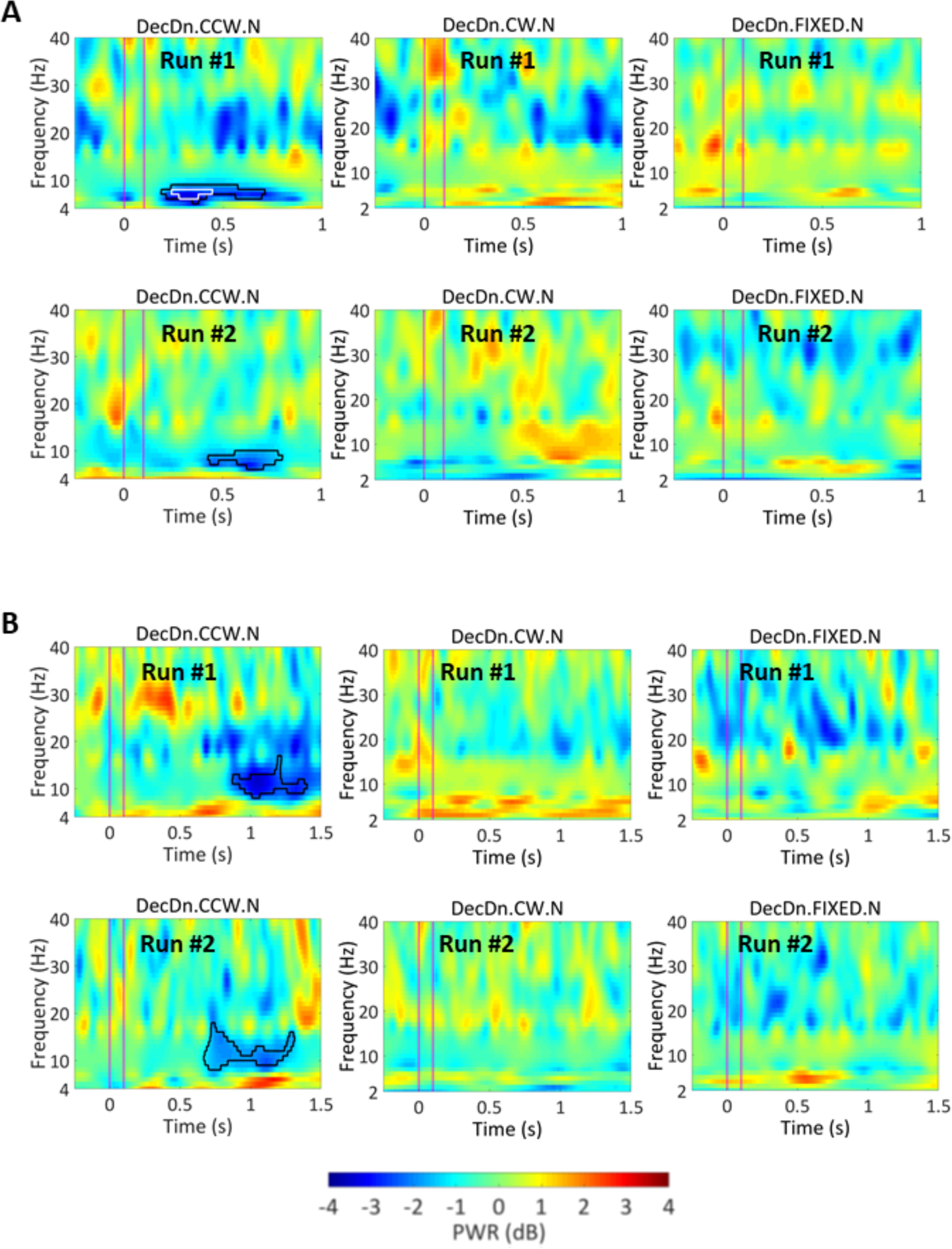
Repeated results from two strongly-responding participants. In both (A) and (B), participants were tested weeks or months apart under the same conditions (Run #1 and Run #2). Time/frequency maps show similar timing and bandwidth of significant alpha power changes (blue clusters in outlines) after counterclockwise rotation, while activity outside the alpha-ERD response, and activity in other conditions is inconsistent across runs. Black/white outlines indicate significance at the p<0.05 and p<0.01 thresholds. The participant in (A) had an alpha peak frequency at <9 Hz and a lower-frequency alpha-ERD response. The participant in (B) had an alpha peak frequency >11 Hz and a higher-frequency alpha-ERD response. Minor power fluctuations in the other conditions or in different frequency bands were not repeated across runs, indicating that only the alpha-ERD was a repeatable signature of magnetosensory processing.

**Video 1.** Test of the electrical induction mechanism of magnetoreception using data from a participant with a strong, repeatable alpha-ERD magnetosensory response. Bottom row shows the DecDn.CCW.N, DecDn.CW.N and DecDn.FIXED.N conditions (64 trials per condition) of the DecDn.N experiment; top row shows the corresponding conditions for the DecUp.N experiment. Scalp topography changes from −0.25 s pre-stimulus to +1 s post-stimulus. The CCW rotation of a downwards-directed field (DecDn.CCW.N) caused a strong, repeatable alpha-ERD (lower left panel, p<0.01 at Fz); weak alpha power fluctuations observed in other conditions (DecDn.CW.N, DecDn.FIXED.N, DecUp.CW.N, DecUp.CCW.N and DecUp.FIXED.N) were not consistent across multiple runs of the same experiment. If the magnetoreception mechanism is based on electrical induction, the same response should occur in conditions with identical ∂**B**/∂**t**(DecDn.CCW.N and DecUp.CCW.N), but the response was observed only in one of these conditions: a result that contradicts the predictions of the electrical induction hypothesis.

**Video 2.** Test of the quantum compass mechanism of magnetoreception using data from another strongly-responding participant. Bottom vs. top rows compare the DecDn.N and DecUp.S experiments in the CCW, CW and FIXED conditions (DecDn.CCW.N, DecDn.CW.N, DecDn.FIXED.N, DecUp.CW.S, DecUp.CCW.S and DecUp.FIXED.S with 100 trials per condition). The quantum compass is not sensitive to magnetic field polarity, so magnetosensory responses should be identical for the DecDn.CCW.N and DecUp.CCW.S rotations sharing the same axis. Our results contradict this prediction. A significant, repeatable alpha-ERD is only observed in the DecDn.CCW.N condition (lower left panel, p<0.01 at Fz), with no strong, consistent effects in the DecUp.CCW.S condition (top left panel) or any other condition.

### Extended Discussion

#### 6. Controlling for Magnetomechanical Artifacts

A question that arises in all studies of human perception is whether confounding artifacts in the experimental system produced the observed effects. The Sham experiments using double-wrapped, bonded coil systems controlled by remote computers and power supplies indicate that obvious artifacts such as resistive warming of the wires or magnetomechanical vibrations between adjacent wires are not responsible. In Active mode, however, magnetic fields produced by the coils interact with each other with maximum torques occurring when the moment **u** of one coil set is orthogonal to the field **B** of another (torque = **u** × **B**). Hence, small torques on the coils might produce transient, sub-aural motion cues. Participants might detect these cues subconsciously even though the coils are anchored to the Faraday cage at many points; the chair and floor assemblies are mechanically isolated from the coils; the experiments are run in total darkness, and the effective frequencies of change are all below 5 Hz and acting for only 0.1 second. No experimenters or participants ever claimed to perceive field rotations consciously even when the cage was illuminated and efforts were made to consciously detect the field rotations. Furthermore, the symmetry of the field rotations and the asymmetric nature of the results both argue strongly against this type of artifact. During the declination experiments, for example, the vertical component of the magnetic field is held constant while a constant-magnitude horizontal component is rotated 90° via the N/S and E/W coil axes. Hence, the torque pattern produced by DecDn.CCW.N rotations should be identical to that of the DecUp.CW.S rotations, yet these conditions yielded dramatically different results. We conclude that magnetomechanical artifacts are not responsible for the observed responses.

#### 7. Testing for Artifacts or Perception from Electrical Induction

Another source of artifacts might be electrical eddy currents induced during field sweeps that might stimulate subsequent EEG brain activity in the head or perhaps in the skin or scalp adjacent to EEG sensors. Such artifacts would be hard to distinguish from a magnetoreceptive structure based on electrical induction. For example, the alpha-ERD effects might arise via some form of voltage-sensitive receptor in the scalp subconsciously activating sensory neurons and transmitting information to the brain for further processing. However, for any such electrical induction mechanism the Maxwell-Faraday law holds that the induced electric field **E** is related to the magnetic field vector, **B**(t), by:

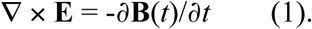

During a declination rotation, the field vector **B**(*t*) is given by: **B**(*t*) = **B**_V_ + **B**_H_(*t*), where **B**_V_ is the constant vertical field component, *t* is time, **B**_H_(t) is the rotating horizontal component, and the quantities in **bold** are vectors. Because the derivative of a constant is zero, the static vertical vector **B**_V_ has no effect, and the induced electrical effect depends only on the horizontally-rotating vector, **B**_H_(t):

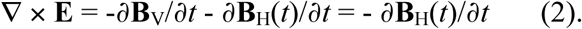

As noted in the main text, Video 1 shows results for the induction test shown in Fig. 2C for which the sweeps of the horizontal component are identical, going along a 90° arc between NE and NW (DecDn.CCW.N and DecUp.CCW.N). The two trials differ only by the direction of the static vertical vector, **B**_V_, which is held in the downwards orientation for the bottom row of Video 1 and upwards in the top row. As only the DecDn.CCW.N sweep elicits alpha-ERD, and the DecUp.CCW.N sweep does not elicit alpha-ERD, electrical induction cannot be the mechanism for this effect either via some artifact of the EEG electrodes or an intrinsic anatomical structure.

We also ran additional control experiments on “EEG phantoms,” which allow us to isolate the contribution of environmental noise and equipment artifacts. Typical setups range from simple resistor circuits to fresh human cadavers. We performed measurements on two commonly-used EEG phantoms: a bucket of saline, and a cantaloupe. From these controls, we isolated the electrical effects induced by magnetic field rotations. The induced effects were similar to the artifact observed in human participants during the 100 ms magnetic stimulation interval. In cantaloupe and in the water-bucket controls, no alpha-ERD responses were observed in Active or Sham modes suggesting that a brain is required to produce a magnetosensory response downstream of any induction artifacts in the EEG signal.

#### 8. Non-polar magnetoreceptivity (attributed to birds) cannot explain the present data

Birds and some other animals display a magnetic inclination compass that identifies the steepest angle of magnetic field dip with respect to gravity (R. Wiltschko & W. Wiltschko, 1995; W. Wiltschko, 1972), and as noted earlier this compass can operate at dips as shallow as 5° from horizontal (Schwarze et al., 2016). This allows a bird to identify the direction of the closest pole (North or South) without knowing the polarity of the magnetic field. If a bird knows it is in the (e.g.) Northern Hemisphere, it can use this maximum dip to identify the direction of geographic North. However, this mechanism could not distinguish between the antipodal (vector opposite) fields used in our biophysical test of polarity sensitivity. If we create a field with magnetic north down and to the front, the bird would correctly identify North as forward. However, if we point magnetic north up and to the back, the bird would still identify North as forward because that is the direction of maximum dip.

Because magnetism and gravity are distinct, non-interacting forces of nature, the observed behavior must arise from neural processing of sensory information from separate transduction mechanisms (J. L. Kirschvink et al., 2010). If polarity information is not present initially from a magnetic transducer or lost in subsequent neural processing, it cannot be recovered by adding information from other sensory modalities. As an illustration, if we gave our participants a compass with a needle that did not have its North tip marked, they could not distinguish the polarity of an applied magnetic field even if we gave them a gravity pendulum or any other non-magnetic sensor.

At present our experimental results in humans suggest the combination of a magnetic and a positional cue. However, we cannot tell if this positional cue is a reference frame using gravity or one aligned with respect to the human body. This could perhaps be addressed by modifying the test chamber to allow the participant to rest in different orientations with respect to gravity.

#### 9. Sastre *et al.* EEG Study

Our results perhaps shed light on a previous study attempting to detect the presence of a low-frequency magnetic stimulus on human brainwaves, which found no significant effects. As part of a major initiative to investigate possible electromagnetic effects on cancer by the US National Institute of Health and the Department of Energy during the 1990’s, Sastre *et al.* (Sastre et al., 2002) analyzed EEG signals for power changes in several frequency bands averaged over 4 s intervals before and after changes in the background magnetic field. However, they did not do the time/frequency analysis that we report here nor averaging of repeated rotations over many trials; wavelet methods were not used as frequently at that time. To test the impact of these differences in data analysis algorithms, we analyzed our data using the techniques in Sastre *et al.* These analyses did not reveal any significant differences in total or band-specific power between any conditions. Thus, our results are consistent with previous findings.

Other differences between our studies lie in the stimulation parameters. In four of seven conditions from Sastre *et al.* (A, B, C and D), the field intensities used (90 μT) were twice as strong as the ambient magnetic field in Kansas City (45 μT) and were well above intensity alterations known to cause birds to ignore geomagnetic cues (W. Wiltschko, 1972). Additionally, Sastre *et al.* chose to use clockwise but not counterclockwise rotations (conditions B and C). In our study, rotating the declination clockwise did not yield statistically significant effects although the reasons are not yet understood (Table 1).

## Acknowledgements

This work was supported directly by Human Frontiers Science Program grant HFSP-RGP0054/2014 to S.S., J.L.K. and A.M., and more recent analysis of data was supported by DARPA RadioBio Program grant (D17AC00019) to JLK and SS, and Japan Society for the Promotion of Science (JSPS) KAKENHI grant 18H03500 to AM. Previous support to J.L.K. from the Fetzer institute allowed construction of an earlier version of the 2 m Merritt coil system. C.X.W. and S.S. have been partly supported by JST.CREST. We thank Dragos Harabor, James Martin, Kristján Jónsson, Mara Green and Sarah Crucilla for work on earlier versions of this project and other members of the Kirschvink, Shimojo, and Matani labs for discussions and suggestions. We also thank James Randi, co-founder of the Committee for the Scientific Investigation of Claims of the Paranormal (CSICOP), for advice on minimizing potential artifacts in the experimental design. Dr. Heinrich Mouritsen of the University of Oldenberg gave valuable advice for construction of the Faraday cage and input on an earlier draft of the manuscript.

## Author Contributions

J.L.K. initiated, and with S.S. and A.M., planned and directed the research. C.X.W., D.A.W. and I.A.H. largely designed the stimulation protocols and conducted the experiments and data analysis. C.P.C., J.N.H.A., S.E.B. and Y.M. designed and built the Faraday cage and implemented the magnetic stimulation protocols. All authors contributed to writing and editing the manuscript.

## Online Content

All digital data are available at https://data.caltech.edu/records/930 and https://data.caltech.edu/records/931, including MatLab^™^ scripts used for the automatic data analysis.

## References

Able, K. P., & Gergits, W. F. (1985). Human Navigation: Attempts to Replicate Baker’s Displacement Experiment. In J. L. Kirschvink, D. S. Jones, & B. J. MacFadden. (Eds.), Magnetite Biomineralization and Magnetoreception in Organisms: A New Biomagnetism (pp. 569–572). New York: Plenum Press.

Baker, R. R. (1980). Goal orientation by blindfolded humans after long-distance displacement: possible involvement of a magnetic sense. Science, 210(4469), 555–557.

Baker, R. R. (1982). Human navigation and the 6th sense: Simon and Schuster.

Baker, R. R. (1987). Human Navigation and Magnetoreception - the Manchester Experiments Do Replicate. Animal Behaviour, 35, 691–704. doi:Doi 10.1016/S0003- 3472(87)80105–7

Bazylinski, D. A., Schlezinger, D. R., Howes, B. H., Frankel, R. B., & Epstein, S. S. (2000). Occurrence and distribution of diverse populations of magnetic protists in a chemically stratified coastal salt pond. Chemical Geology, 169(3–4), 319–328. doi:Doi 10.1016/S0009–2541(00)00211–4

Beason, R. C., & Semm, P. (1996). Does the avian ophthalmic nerve carry magnetic navigational information? Journal of Experimental Biology, 199(5), 1241–1244.

Beason, R. C., Wiltschko, R., & Wiltschko, W. (1997). Pigeon homing: Effects of magnetic pulses on initial orientation. Auk, 114(3), 405–415. doi:Doi 10.2307/4089242

Benjamini, Y., & Hochberg, Y. (1995). Controlling the False Discovery Rate - a Practical and Powerful Approach to Multiple Testing. Journal of the Royal Statistical Society Series B-Methodological, 57(1), 289–300.

Block, S. M. (1992). Biophysical Aspects of Sensory Transduction. In D. P. Corey & S. D. Roper (Eds.), Sensory Transduction (Vol. 45, pp. 424). Marine Biological Laboratory, Woods Hole, Massachusetts: Rockfeller University Press.

Boorman, G. A., Bernheim, N. J., Galvin, M. J., Newton, S. A., Parham, F. M., Portier, C. J., & Wolfe, M. S. (1999). NIEHS Report on Health Effects from Exposure to Power-Line FrequeNCy Electric and Magnetic Fields (NIEHS Ed. Vol. 99–4493). Research Triangle Park, NC 27709: Department of Health & Humn Services, US Government.

Carporzen, L., Weiss, B. P., Gilder, S. A., Pommier, A., & Hart, R. J. (2012). Lightning remagnetization of the Vredefort impact crater: No evidence for impact-generated magnetic fields. Journal of Geophysical Research-Planets, 117. doi:Artn E01007 10.1029/2011je003919

Cohen, M. X. (2014). Analyzing Neural Time Series Data Theory and Practice Preface. Cambridge, Massachusetts: MIT Press.

Counselman, C. (2013). Excellent, Easy, Cheap Common-Mode Chokes. National Contest Journal of the American Radio Relay League, 41(1), 3–5.

Diaz Ricci, J. C., & Kirschvink, J. L. (1992). Magnetic domain state and coercivity predictions for biogenic greigite (Fe3S4): A comparison of theory with magnetosome observations. J. Geophys. Res., 97, 17309–17315.

Diebel, C. E., Proksch, R., Green, C. R., Neilson, P., & Walker, M. M. (2000). Magnetite defines a vertebrate magnetoreceptor. Nature, 406(6793), 299–302. doi:10.1038/35018561

Dunn, J. R., Fuller, M., Zoeger, J., Dobson, J., Heller, F., Hammann, J., … Moskowitz, B. M. (1995). Magnetic material in the human hippocampus. Brain Res Bull, 36(2), 149–153.

Eder, S. H., Cadiou, H., Muhamad, A., McNaughton, P. A., Kirschvink, J. L., & Winklhofer, M. (2012). Magnetic characterization of isolated candidate vertebrate magnetoreceptor cells. Proc Natl Acad Sci U S A, 109(30), 12022–12027. doi:10.1073/pnas.1205653109

Elbers, D., Bulte, M., Bairlein, F., Mouritsen, H., & Heyers, D. (2017). Magnetic activation in the brain of the migratory northern wheatear (Oenanthe oenanthe). J Comp Physiol A Neuroethol Sens Neural Behav Physiol, 203(8), 591–600. doi:10.1007/s00359–017–1167–7

Engels, S., Schneider, N. L., Lefeldt, N., Hein, C. M., Zapka, M., Michalik, A., … Mouritsen, H. (2014). Anthropogenic electromagnetic noise disrupts magnetic compass orientation in a migratory bird. Nature, 509(7500), 353–356. doi:10.1038/nature13290

Ernst, D. A., & Lohmann, K. J. (2016). Effect of magnetic pulses on Caribbean spiny lobsters: implications for magnetoreception. J Exp Biol, 219(Pt 12), 1827–1832. doi:10.1242/jeb.136036

Fillmore, E. P., & Seifert, M. F. (2015). Anatomy of the Trigeminal Nerve. In R. S. Tubbs, E. Rizk, M. Shoja, M. Loukas, N. Barbaro, & R. Spinner (Eds.), Nerves and Nerve Injuries (Vol. 1, pp. 319–350): Academic Press.

Frankel, R. B., & Blakemore, R. P. (1980). Navigational Compass in Magnetic Bacteria. Journal of Magnetism and Magnetic Materials, 15–8(Jan-), 1562–1564. doi:Doi 10.1016/0304–8853(80)90409–6

Gilder, S. A., Wack, M., Kaub, L., Roud, S. C., Petersen, N., Heinsen, H., … Schmitz, C. (2018). Distribution of magnetic remanence carriers in the human brain. Scientific Reports, 8(11363). doi:DOI:10.1038/s41598–018–29766-z 1

Gould, J. S., & Able, K. P. (1981). Human homing: an elusive phenomenon. Science, 212(4498), 1061–1063.

Hand, E. (2016). Polar explorer. Science, 352(6293), 1508–1510, 1512–1503. doi:10.1126/science.352.6293.1508

Hartmann, T., Schlee, W., & Weisz, N. (2012). It’s only in your head: expectancy of aversive auditory stimulation modulates stimulus-induced auditory cortical alpha desynchronization. Neuroimage, 60(1), 170–178. doi:10.1016/j. neuroimage.2011.12.034

Haviland, J. B. (1998). Guugu Yimithir cardinal directions. Ethos, 26(1), 25–47. doi:DOI 10.1525/eth.1998.26.1.25

Holland, R. A. (2010). Differential effects of magnetic pulses on the orientation of naturally migrating birds. J R Soc Interface, 7(52), 1617–1625. doi:10.1098/rsif.2010.0159

Holland, R. A., & Helm, B. (2013). A strong magnetic pulse affects the precision of departure direction of naturally migrating adult but not juvenile birds. Journal of the Royal Society Interface, 10, 20121047.

Holland, R. A., Kirschvink, J. L., Doak, T. G., & Wikelski, M. (2008). Bats use magnetite to detect the earth’s magnetic field. Plos One, 3(2), e1676. doi:10.1371/journal.pone.0001676

Hore, P. J., & Mouritsen, H. (2016). The Radical-Pair Mechanism of Magnetoreception. In K. A. Dill (Ed.), Annual Review of Biophysics, Vol 45 (Vol. 45, pp. 299–344).

Irwin, W. P., & Lohmann, K. J. (2005). Disruption of magnetic orientation in hatchling loggerhead sea turtles by pulsed magnetic fields. J Comp Physiol A Neuroethol Sens Neural Behav Physiol, 191(5), 475–480. doi:10.1007/s00359–005–0609–9

Johnsen, S., & Lohmann, K. J. (2008). Magnetoreception in animals. Physics Today, 61(3), 2935. doi:Doi 10.1063/1.2897947

Kalmijn, A. J. (1981). Biophysics of Geomagnetic-Field Detection. IEEE Transactions on Magnetics, 17(1), 1113–1124. doi:Doi 10.1109/Tmag.1981.1061156

Kirschvink, J., Padmanabha, S., Boyce, C., & Oglesby, J. (1997). Measurement of the threshold sensitivity of honeybees to weak, extremely low-frequency magnetic fields. J Exp Biol, 200(Pt 9), 1363–1368.

Kirschvink, J. L. (1992). Uniform magnetic fields and Double-wrapped coil systems: Improved techniques for the design of biomagnetic experiments. Bioelectromagnetics, 13, 401–411.

Kirschvink, J. L., & Gould, J. L. (1981). Biogenic magnetite as a basis for magnetic field sensitivity in animals. Bio Systems, 13, 181–201.

Kirschvink, J. L., & Kobayashi-Kirschvink, A. (1991). Is geomagnetic sensitivity real? Replication of the Walker-Bitterman conditioning experiment in honey bees. American Zoologist, 31, 169–185.

Kirschvink, J. L., Kobayashi-Kirschvink, A., & Woodford, B. J. (1992). Magnetite biomineralization in the human brain. Proc Natl Acad Sci U S A, 89(16), 7683–7687.

Kirschvink, J. L., Winklhofer, M., & Walker, M. M. (2010). Biophysics of magnetic orientation: strengthening the interface between theory and experimental design. J R Soc Interface, 7 Suppl 2, S179–191. doi:10.1098/rsif.2009.0491.focus

Klimesch, W. (1999). EEG alpha and theta oscillations reflect cognitive and memory performance: a review and analysis. Brain Res Brain Res Rev, 29(2–3), 169–195.

Klimesch, W., Doppelmayr, M., Russegger, H., Pachinger, T., & Schwaiger, J. (1998). Induced alpha band power changes in the human EEG and attention. Neurosci Lett, 244(2), 73–76. doi:10.1016/s0304–3940(98)00122–0

Kobayashi, A., & Kirschvink, J. L. (1995). Magnetoreception and EMF Effects: Sensory Perception of the geomagnetic field in Animals & Humans. In M. Blank (Ed.), Electromagnetic Fields: Biological Interactions and Mechanisms (pp. 367–394). Washington, DC.: American Chemical Society Books.

Kramer, G. (1953). Wird die Sonnenhohe bei der Heimfindeorientierung verwertet? Journal of Ornitholgy, 94, 201–219.

Landler, L., Painter, M. S., Youmans, P. W., Hopkins, W. A., & Phillips, J. B. (2015). Spontaneous magnetic alignment by yearling snapping turtles: rapid association of radio frequency dependent pattern of magnetic input with novel surroundings. Plos One, 10(5), e0124728. doi:10.1371/journal.pone.0124728

Levinson, S. C. (2003). Space in Language and Cognition. Cambridge: Cambridge University Press.

Light, P., Salmon, M., & Lohmann, K. J. (1993). Geomagnetic Orientation of Loggerhead Sea-Turtles – Evidence for an Inclination Compass. Journal of Experimental Biology, 182, 1–9.

Liu, G. T. (2005). The trigeminal nerve and its central connections. In N. R. Miller, N. J. Newman, V. Biousse, & K. J. B. (Eds.), Walsh & Hoyt’s Clinical Neuro-Ophthalmalogy, 6th edition (6th ed., Vol. 1, pp. 1233–1268). Philadelphia: Lippencott Williams & Wilkins.

Lohmann, K. J., Cain, S. D., Dodge, S. A., & Lohmann, C. M. (2001). Regional magnetic fields as navigational markers for sea turtles. Science, 294(5541), 364–366. doi:10.1126/science.1064557

Lohmann, K. J., & Lohmann, C. M. F. (1996). Detection of magnetic field intensity by sea turtles. Nature, 380(6569), 59–61. doi:DOI 10.1038/380059a0

Lopez, C., & Blanke, O. (2011). The thalamocortical vestibular system in animals and humans. Brain Res Rev, 67(1–2), 119–146. doi:10.1016/j. brainresrev.2010.12.002

Maher, B. A., Ahmed, I. A., Karloukovski, V., MacLaren, D. A., Foulds, P. G., Allsop, D., … Calderon-Garciduenas, L. (2016). Magnetite pollution nanoparticles in the human brain. Proc Natl Acad Sci U S A, 113(39), 10797–10801. doi:10.1073/pnas.1605941113

Martin, H., & Lindauer, M. (1977). Der Einfluss der Erdmagnetfelds und die Schwerorientierung der Honigbiene. J. Comp. Physiol., 122,, 145–187.

Meakins, F. (2011). Spaced Out: Intergenerational Changes in the Expression of Spatial Relations by Gurindji People. Australian Journal of Linguistics, 31(1), 43–77. doi:Pii 932432693 10.1080/07268602.2011.532857

Meakins, F., & Algy, C. (2016). Deadly Reckoning: Changes in Gurindji Children’s Knowledge of Cardinals. Australian Journal of Linguistics, 36(4), 479–501. doi:10.1080/07268602.2016.1169973

Meakins, F., Jones, C., & Algy, C. (2016). Bilingualism, language shift and the corresponding expansion of spatial cognitive systems. Language Sciences, 54, 1–13. doi:10.1016/j. langsci.2015.06.002

Merritt, R., Purcell, C., & Stroink, G. (1983). Uniform Magnetic-Field Produced by 3-Square, 4-Square, and 5-Square Coils. Review of Scientific Instruments, 54(7), 879–882. doi:Doi 10.1063/1.1137480

Mora, C. V., Davison, M., Wild, J. M., & Walker, M. M. (2004). Magnetoreception and its trigeminal mediation in the homing pigeon. Nature, 432(7016), 508–511. doi:10.1038/nature03077

Munro, U., Munro, J. A., Phillips, J. B., Wiltschko, R., & Wiltschko, W. (1997). Evidence for a magnetite-based navigational ’’map’’ in birds. Naturwissenschaften, 84(1), 26–28. doi:DOI 10.1007/s001140050343

Munro, U., Munro, J. A., Phillips, J. B., & Wiltschko, W. (1997). Effect of wavelength of light and pulse magnetisation on different magnetoreception systems in a migratory bird. Australian Journal of Zoology, 45(2), 189–198. doi:Doi 10.1071/Zo96066

Nuwer, M. R., Comi, G., Emerson, R., Fuglsang-Frederiksen, A., Guerit, J. M., Hinrichs, H., … Rappelsburger, P. (1998). IFCN standards for digital recording of clinical EEG. International Federation of Clinical Neurophysiology. Electroencephalogr Clin Neurophysiol, 106(3), 259–261. doi:10.1016/s0013–4694(97)00106–5

Peng, W., Hu, L., Zhang, Z., & Hu, Y. (2012). Causality in the association between P300 and alpha event-related desynchronization. Plos One, 7(4), e34163. doi:10.1371/journal.pone.0034163

Pfurtscheller, G., & Lopes da Silva, F. H. (1999). Event-related EEG/MEG synchronization and desynchronization: basic principles. Clin Neurophysiol, 110(11), 1842–1857. doi:10.1016/s1388–2457(99)00141–8

Pfurtscheller, G., Neuper, C., & Mohl, W. (1994). Event-related desynchronization (ERD) during visual processing. Int J Psychophysiol, 16(2–3), 147–153. doi:10.1016/0167- 8760(89)90041-x

Poisson, R. J., & Miller, M. E. (2014). Spatial disorientation mishap trends in the U.S. Air force 1993–2013. Aviat Space Environ Med, 85(9), 919–924. doi:10.3357/ASEM.3971.2014

Ritz, T., Adem, S., & Schulten, K. (2000). A model for photoreceptor-based magnetoreception in birds. BiophysJ, 78(2), 707–718. doi:10.1016/S0006- 3495(00)76629-X

Rosenblum, B., Jungerman, R. L., & Longfellow, L. (1985). Limits to induction-based magnetoreception. In J. L. Kirschvink, D. S. Jones, & B. J. MacFadden (Eds.), Magnetite Biomineralization and Magnetoreception in Organisms: A New Biomagnetism (pp. 223–232). New York: Plenum Press.

Saper, C. B. (2002). The central autonomic nervous system: conscious visceral perception and autonomic pattern generation. Annu Rev Neurosci, 25, 433–469. doi:10.1146/annurev.neuro.25.032502.111311

Sastre, A., Graham, C., Cook, M. R., Gerkovich, M. M., & Gailey, P. (2002). Human EEG responses to controlled alterations of the Earth’s magnetic field. Clin Neurophysiol, 113(9), 1382–1390. doi:10.1016/s1388–2457(02)00186–4

Schulten, K. (1982). Magnetic-Field Effects in Chemistry and Biology. Festkorperprobleme-Advances in Solid State Phyics, 22, 61–83. doi:DOI 10.1007/BFb0107935

Schultheiss-Grassi, P. P., Wessiken, R., & Dobson, J. (1999). TEM investigations of biogenic magnetite extracted from the human hippocampus. Biochim Biophys Acta, 1426(1), 212–216. doi:10.1016/s0304–4165(98)00160–3

Schwarze, S., Steenken, F., Thiele, N., Kobylkov, D., Lefeldt, N., Dreyer, D., … Mouritsen, H. (2016). Migratory blackcaps can use their magnetic compass at 5 degrees inclination, but are completely random at 0 degrees inclination. Sci Rep, 6, 33805. doi:10.1038/srep33805

Semm, P., & Beason, R. C. (1990). Responses to small magnetic variations by the trigeminal system of the bobolink. Brain Res Bull, 25(5), 735–740. doi:10.1016/0361- 9230(90)90051-z

Tomanova, K., & Vacha, M. (2016). The magnetic orientation of the Antarctic amphipod Gondogeneia antarctica is cancelled by very weak radiofrequency fields. J Exp Biol, 219(Pt 11), 1717–1724. doi:10.1242/jeb.132878

Tukey, J. W. (1949). Comparing individual means in the analysis of variance. Biometrics, 5(2), 99–114. doi:10.2307/3001913

Veniero, D., Bortoletto, M., & Miniussi, C. (2009). TMS-EEG co-registration: On TMS-induced artifact. Clinical Neurophysiology, 120(7), 1392–1399. doi:10.1016/j.clinph.2009.04.023

Walker, M. M., Dennis, T. E., & Kirschvink, J. L. (2002). The magnetic sense and its use in long-distance navigation by animals. Current Opinion in Neurobiology, 12(6), 735–744. doi:Doi 10.1016/S0959–4388(02)00389–6

Walker, M. M., Diebel, C. E., Haugh, C. V., Pankhurst, P. M., Montgomery, J. C., & Green, C. R. (1997). Structure and function of the vertebrate magnetic sense. Nature, 390(6658), 371–376.

Wegner, R. E., Begall, S., & Burda, H. (2006). Magnetic compass in the cornea: local anaesthesia impairs orientation in a mammal. J Exp Biol, 209(Pt 23), 4747–4750. doi:10.1242/jeb.02573

Welch, P. D. (1967). Use of Fast Fourier Transform for Estimation of Power Spectra - a Method Based on Time Averaging over Short Modified Periodograms. Ieee Transactions on Audio and Electroacoustics, Au15(2), 70-+. doi:Doi 10.1109/Tau.1967.1161901

Westby, G. W. M., & Partridge, K. J. (1986). Human Homing - Still No Evidence Despite Geomagnetic Controls. Journal of Experimental Biology, 120, 325–331.

Wiltschko, R., Thalau, P., Gehring, D., Niessner, C., Ritz, T., & Wiltschko, W. (2015). Magnetoreception in birds: the effect of radio-frequency fields. J R Soc Interface, 12(103). doi:10.1098/rsif.2014.1103

Wiltschko, R., & Wiltschko, W. (1995). Magnetic orientation in animals (Vol. 33). Berlin: Springer.

Wiltschko, W. (1972). The influence of magnetic total intensity and inclination on directions preferred by migrating European robins (Erithacus rubeccula). In S. R. Galler, K. Schmidt-Koenig, G. J. Jacobs, & R. E. Belleville (Eds.), Animal Orientation and Navigation (Vol. NASA SP-262, pp. 569–578). Washington, D.C., USA: U.S. Government Printing Office.

Wiltschko, W., Ford, H., Munro, U., Winklhofer, M., & Wiltschko, R. (2007). Magnetite-based magnetoreception: the effect of repeated pulsing on the orientation of migratory birds. J Comp Physiol A Neuroethol Sens Neural Behav Physiol, 193(5), 515–522. doi:10.1007/s00359–006–0207–5

Wiltschko, W., Munro, U., Beason, R. C., Ford, H., & Wiltschko, R. (1994). A Magnetic Pulse Leads to a Temporary Deflection in the Orientation of Migratory Birds. Experientia, 50(7), 697–700. doi:Doi 10.1007/Bf01952877

Wiltschko, W., Munro, U., Ford, H., & Wiltschko, R. (1998). Effect of a magnetic pulse on the orientation of silvereyes, zosterops l. lateralis, during spring migration. J Exp Biol, 201 (Pt 23)(23), 3257–3261.

Wiltschko, W., Munro, U., Ford, H., & Wiltschko, R. (2009). Avian orientation: the pulse effect is mediated by the magnetite receptors in the upper beak. Proc Biol Sci, 276(1665), 2227–2232. doi:10.1098/rspb.2009.0050

Wiltschko, W., Munro, U., Wiltschko, R., & Kirschvink, J. L. (2002). Magnetite-based magnetoreception in birds: the effect of a biasing field and a pulse on migratory behavior. J Exp Biol, 205(Pt 19), 3031–3037.

Wiltschko, W., & Wiltschko, R. (1995). Migratory Orientation of European Robins Is Affected by the Wavelength of Light as Well as by a Magnetic Pulse. Journal of Comparative Physiology a-Sensory Neural and Behavioral Physiology, 1 77(3), 363–369.

Yeagley, H. L. (1947). A Preliminary Study of a Physical Basis of Bird Navigation. Journal of Applied Physics, 18(12), 1035–1063. doi:Doi 10.1063/1.1697587

